# Engineering CAR-NK cells to secrete IL15 sustains their anti-AML functionality, but is associated with systemic toxicities

**DOI:** 10.1101/2021.09.23.461509

**Authors:** Ilias Christodoulou, Won Jin Ho, Andrew Marple, Jonas W. Ravich, Ada Tam, Ruyan Rahnama, Adam Fearnow, Cambrynne Rietberg, Sean Yanik, Elena E. Solomou, Ravi Varadhan, Michael A. Koldobskiy, Challice L. Bonifant

**Author notes:** **Corresponding author:** Challice Bonifant, MD, PhD.

## Abstract

**Background:** The prognosis of patients with recurrent/refractory acute myelogenous leukemia (AML) remains poor and cell-based immunotherapies hold promise to improve outcomes. NK cells can elicit an anti-leukemic response via a repertoire of activating receptors that bind AML surface ligands. NK cell adoptive transfer is safe but thus far has shown limited anti-AML efficacy. Here, we aimed to overcome this limitation by engineering NK cells to express chimeric antigen receptors (CARs) to boost their anti-AML activity, and interleukin-15 (IL15) to enhance their persistence.

**Methods:** We characterized in detail NK cell populations expressing a panel of AML (CD123)-specific CARs and/or IL15 *in vitro* and in AML xenograft models.

**Results:** CARs with 2B4.ζ or 4-1BB.ζ signaling domains demonstrated greater cell surface expression and endowed NK cells with improved anti-AML activity *in vitro*. Initial *in vivo* testing revealed that only 2B4.ζ CAR-NK cells had improved anti-AML activity in comparison to untransduced (UTD) and 4-1BB.ζ CAR-NK cells. However, the benefit was transient due to limited CAR-NK cell persistence. Transgenic expression of secretory (s)IL15 in 2B4.ζ CAR and UTD NK cells improved their effector function in the setting of chronic antigen simulation *in vitro*. Multiparameter flow analysis after chronic antigen exposure identified the expansion of unique NK cell subsets. 2B4.ζ/sIL15 CAR and sIL15 NK cells maintained an overall activated NK cell phenotype. This was confirmed by transcriptomic analysis, which revealed a highly proliferative and activated signature in these NK cell groups. *In vivo*, 2B4.ζ/sIL15 CAR-NK cells had potent anti-AML activity in one model, while 2B4.ζ/sIL15 CAR and sIL15 NK cells induced lethal toxicity in a second model.

**Conclusion:** Transgenic expression of CD123-CARs and sIL15 enabled NK cells to function in the setting of chronic antigen exposure but was associated with systemic toxicities. Thus, our study provides the impetus to explore inducible and controllable expression systems to provide cytokine signals to AML-specific CAR-NK cells before embarking on early phase clinical testing.

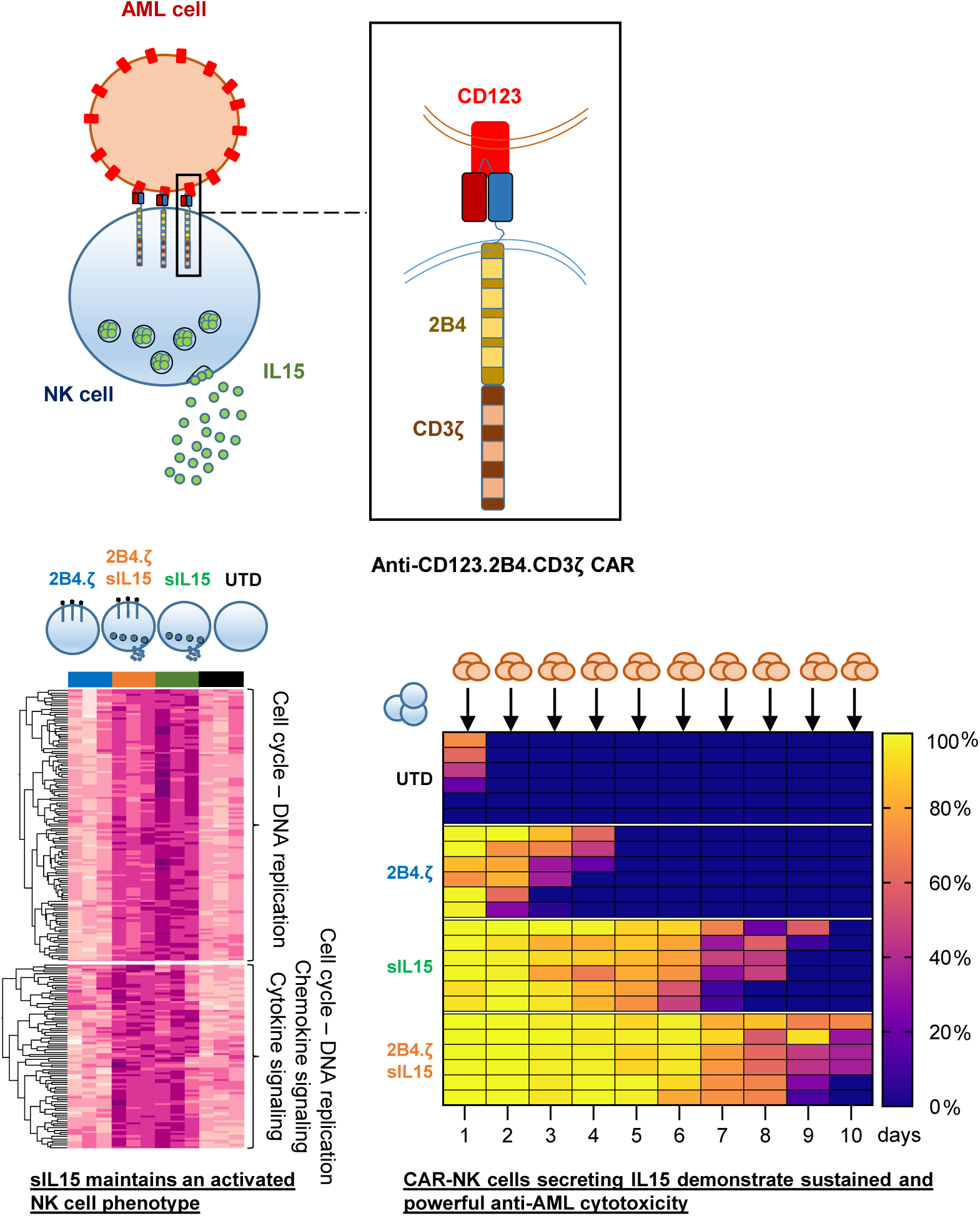

**In Brief:** Secretion of IL15 from anti-CD123.2B4.CD3ζ CAR-NK cells enhances their activation and killing potential against Acute Myelogenous Leukemia, but carries the risk of severe systemic toxicity.

## Introduction

Acute myelogenous leukemia (AML) is a neoplastic disorder characterized by the accumulation of malignant myeloid precursor cells in the bone marrow. AML has an aggressive clinical course in both adults and children.^1,2^ Intensive chemotherapeutic regimens with consolidative hematopoietic cell transplantation (HCT) remain the standard of care, but these treatments can cause significant short and long-term toxicities. Moreover, a subset of patients fail to respond to initial treatment or relapse after chemotherapy +/- HCT. For this reason, targeted therapies with non-overlapping toxicity profiles are aggressively being developed. The alpha chain of the interleukin (IL)-3 receptor (CD123) is highly expressed on both AML blasts and leukemic stem cells,^3,4^ and has been shown to be a safe target in clinical trials of immunotherapeutic agents.^5–7^ Leukemic stem cells are often resistant to chemotherapy and may be most responsible for disease initiation and relapse.^3^ Thus, CD123 targeting therapies could serve as valuable adjunct treatment modalities to achieve and/or sustain remission in high-risk AML patients.

Adoptive cell transfer is a form of anticancer immunotherapy that has promise, and has been successful in the form of CAR-T cell infusions used to treat relapsed/refractory ALL.^8^ CAR-T cells are also being tested against AML in several clinical trials.^9^ CAR-T cell therapies have associated severe toxicities including Cytokine Release Syndrome (CRS), Immune Effector Cell Associated Neurotoxicity Syndrome (ICANS),^10^ and CAR-associated Hemophagocytic Lymphohistiocytosis.^11^ All FDA-approved CAR-T cell therapies originate from autologous hematopoietic starting material. The manufacturing of patient-derived T cell products in the setting of a highly proliferative disease can be challenging.^12,13^ Delay in therapy associated with the time required for per-patient CAR-T manufacturing may not be possible in the setting of uncontrolled AML. In addition, intensive chemotherapy regimens, such as those administered for AML treatment, are associated with poor ex vivo T cell expansion.^14^ Donor T cells can be considered as a product source; however, the infusion of allogeneic T cells is associated with the serious risk of graft-versus-host disease (GVHD).^15^

NK cells are immune effector cells that play a pivotal role as first-line defenders against virally infected or tumor-transformed cells. Though they act with similar cytotoxic mechanisms, NK cell activation and function are distinct from that of T cells. NK cells express a variety of activating receptors (ex. NKG2D, NKp30, NKp46, 2B4 and CD16)^16,17^ that directly interact with cell surface ligands.^16,18^ These activating receptors associate with co-receptors (ex. DAP10, CD3ζ, FcεRIγ) to heighten directed cytotoxicity.^17^ The use of NK cells for immunotherapy has major advantages over the use of T cells. NK cells do not directly cause GVHD, though they may contribute inflammatory mediators that potentiate pre-existing pathology.^15,19^ Thus, they have the potential to be used as an off-the-shelf cellular product that can be manufactured on a large scale and be readily available to patients. This has the advantage of decreasing production costs and preventing manufacturing associated treatment delays. Adoptive NK cell transfer has been shown to be safe in clinical trials, without associated CRS or ICANS.^20^ Because of this safety profile and the expression of NK activating ligands on AML cells,^18^ NK cell therapy has been tested in clinical trials. However, when used as AML treatment, NK cell adoptive transfer induces only transient remission and additional therapies are needed to achieve durable responses.^21–24^

In this study, we engineered NK cells to express CARs with intracellular domains rationally designed as those predicted to enhance NK cell anti-tumor functionality. We demonstrate that NKs expressing CARs targeting CD123 show potent antigen-dependent activation and cytotoxicity. We find CAR expression to be stable and high across a panel of tested receptors incorporating NK-specific activating and costimulatory molecules, with optimal functionality associated with 2B4.ζ and 4-1BB.ζ containing CARs. CAR-NK cells did not demonstrate improved expansion or persistence when compared to unmodified NK cells *in vivo.* We therefore added constitutive IL15 secretion to our CAR-NK cells and observed that this supported an activated phenotype that led to enhanced expansion and anti-tumor cytotoxicity. NK cell activation translated to improved NK cell persistence and expansion *in vivo*. However, constitutive cytokine secretion was also associated with severe toxicity in one animal model. Prior to clinical translation, alternate strategies to activate cytokine signaling in CAR-NK cells should be investigated with a focus to include safety.

## Methods

Details about cell lines, determination of Vector Copy Number (VCN), cytotoxicity assays, cytokine secretion measurement, and RNAseq library preparation and alignment are provided in **Supplemental Methods**.

### Chimeric Antigen Receptor (CAR) Generation

CAR transgenes were designed using the CD123-specific single chain variable fragment (scFv; 26292)^25^ sequence and the hinge, transmembrane, and intracellular domains indicated in Fig. 1A. Sequences were synthesized (GeneArt, ThermoFisher Scientific) and subcloned into pSFG retroviral vectors. All sequences were validated by Sanger Sequencing (Johns Hopkins Genetic Resources Core Facility).

**Figure 1.**
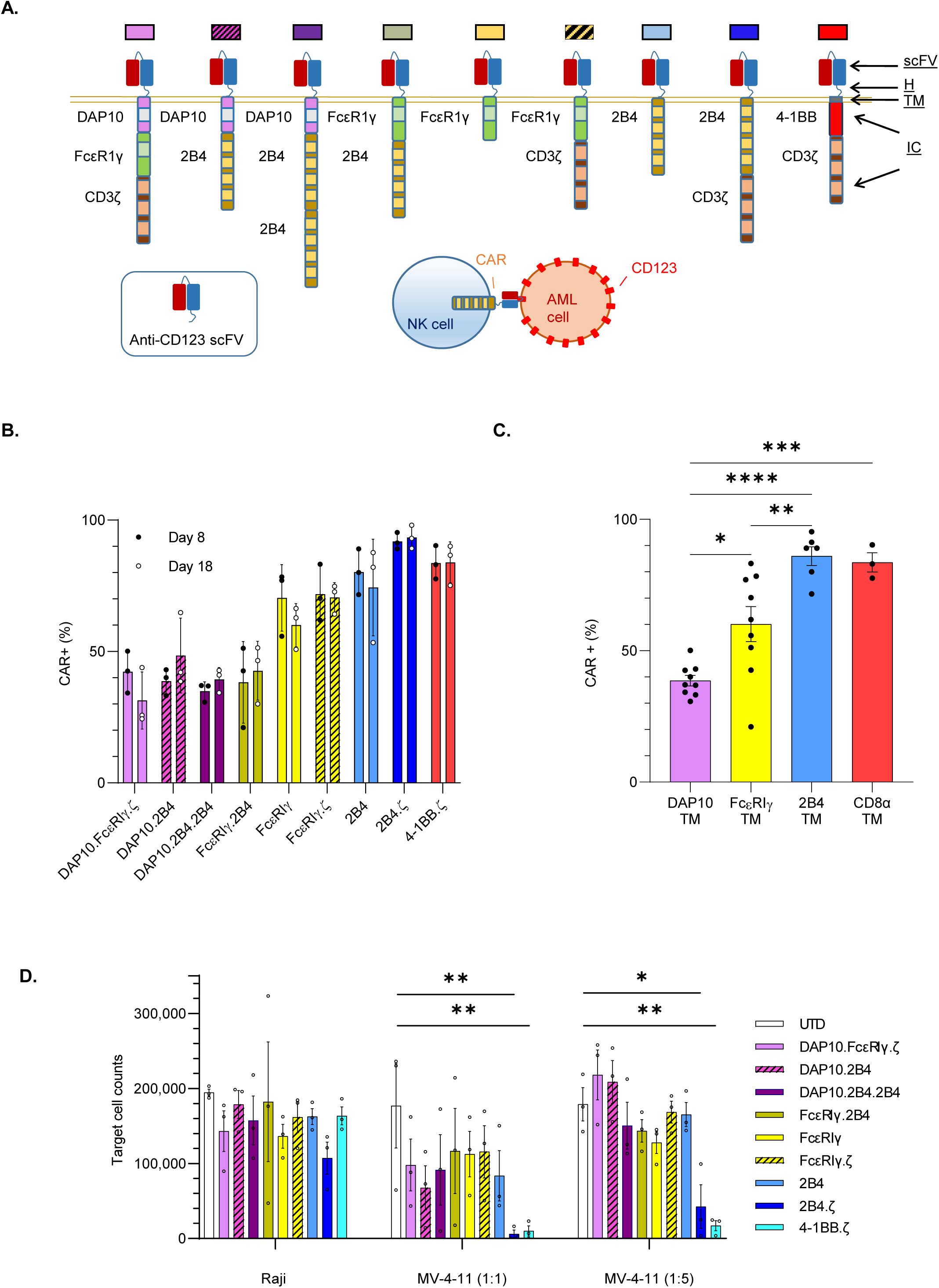
NK cells engineered with anti-CD123 CARs have antigen-specific functionality. (**A**) Schema of CAR design. All CARs bind CD123 via an extracellular single chain variable fragment (scFv). The hinge (H), transmembrane (TM) and intracellular (IC) domains of the CARs are as indicated. Colored boxes represents each particular CAR with colors carried through each figure. (**B**) Percentage (%) of CAR (+) NK cells detected on day (D)8 and D18. (**C**) Bar plot comparing the percentage of CAR (+) NK cells with indicated transmembrane (TM) domains on D8. (**D**) Absolute number of target cells measured after 72h co-culture with indicated NK cells. Initial target cell count was 100,000 in 1:1 and 250,000 in 1:5 E:T ratio conditions. Each bar representative of the mean plus or minus the standard error of mean (+/- SEM); each dot representative of individual NK cell donor. n=3 donors (*p<0.05; **p<0.01; ***p<0.001; ****p<0.0001).

### CAR-NK cell production

Healthy donor peripheral blood mononuclear cells (PBMCs; Anne Arundel Medical Blood Donor Center, Annapolis, MD; Carter Bloodcare, Woodway, TX) were isolated by Ficoll desnity gradient centrifugation and depleted of T cells using CD3-microbeads (Militenyi Biotec, Cologne, Germany). Remaining cells were stimulated on day 0 with lethally irradiated K562 feeder cells^26^ expressing membrane bound IL15 and 4-1BB ligand at a 1:1 ratio. Cells were maintained in SGCM media (CellGenix, Freiburg, Germany) supplemented with 10% Fetal Bovine Serum, 2 mmol/L GlutaMAX (ThermoFisher), and 200 IU/mL hIL-2(Biological Resources Branch Preclinical Biorepository, National Cancer Institute, Frederick, MD). NK cell purity was verified with flow cytometry using fluorophore conjugated antibodies against CD56 and CD3 (Supplemental Table 1). NK cells were transduced on day 4 of culture using transiently produced replication incompetent RD114 pseudotyped retroviral particles immobilized on RetroNectin (Clontech Laborotories, Palo Alto, CA).

### Flow cytometry

Antibodies for NK and cancer cell identification targeted CD56 and CD33 (AML cell lines) or CD19 (Raji) markers. A detailed list of all antibodies, including those used for the evaluation of immune cell phenotype is in Supplemental Table 1. CAR expression analysis was performed using incubation with His-tagged recombinant CD123 protein (SinoBiological, Beijing, China) and secondary staining with αHis-PE or αHis-APC (BioLegend, San Diego, CA). Dead cells were excluded from analysis using LIVE/DEAD Fixable Viability Stains 780 or 575V (BD Horizon, Franklin Lakes, NJ). Cell enumeration was performed with CountBright™ counting beads (ThermoFisher). Human Fc receptors (FcRs) were blocked using Human TruStain FcX™ (BioLegend). In samples stained with multiple BD Horizon Brilliant reagents, Brilliant Stain Buffer Plus (BD Horizon) was used. Compensation was performed with UltraComp eBeads Compensation Beads (ThermoFisher). Cell surface antigens were quantified using microspheres of the Quantum APC Molecules of Equivalent Soluble Fluorochrome (MESF) kit (Bangs Laboratories, Inc.; Fisher, IN). All samples were acquired on FACSCelesta or FACSymphony Cell Analyzers (BD) and analyzed with FlowJo software (v10.6.1; v10.7.2). Cell sorting was performed on FACSMelody (BD).

### Immunophenotype analysis

Two different panels (A and B) were used for evaluation of NK cell receptors and one panel (C) for receptor ligands. Data analysis of the multiparameter panels A and B was performed in R (v3.6.2). The median marker intensities were transformed using arcsinh (inverse hyperbolic sine) with cofactor 150.^27^ Nonlinear dimensionality reduction on randomly selected 500 data points per sample of each panel was performed using uniform manifold approximation and projection (UMAP).^28^ NK cell clusters were identified with the FlowSOM (v1.18.0) algorithm and 40 different metaclusters were generated per panel.^29^ Subsequently, we manually merged hierarchically neighboring clusters similar in biology and median marker intensities. Panel A clusters do not correlate with the ones in panel B.

### Serial Stimulation assay

NK cells were stimulated daily with MV-4-11 cells at a 1:1 effector:target (E:T) ratio in G-rex plates (Wilson Wolf, New Brighton, MN) for a total of ten days. NK cell proliferation and cytotoxicity was measured using flow cytometric analysis. Percent (%) cytotoxicity was calculated based on the target cell numbers on the day of (Y) and the day after (X) stimulation using the formula 100*(X-Y)/(Y). Cell phenotype was evaluated at baseline, on the first (12h) and the tenth (D10) day.

### RNA sequencing

On D10 of serial stimulation, co-culture was depleted first of dead cells using the Dead Cell Removal Kit and next of leukemia cells with CD33-microbeads (Militenyi). NK cell purity was verified with flow cytometry using fluorophore-conjugated CD56 and CD33 antibodies. RNA was extracted from NK cells using the RNeasy Mini Kit (Qiagen, Hilden, Germany) and RNAseq was performed (**Supplemental Methods**). Differential expression analysis and statistical testing was performed using DESeq2 software.^30^

### Xenograft mouse model

All animal studies were carried out under protocols approved by the Johns Hopkins Institutional Animal Care and Use Committee (IACUC). Six to eight week old NSG (NOD.*Cg-Prkdc^scid^Il2rg^tm1Wjl^*/SzJ mice were obtained from an internal colony that originated from the Jackson Laboratory (Bar Harbor, ME). Mice were injected with 1×10^6^ MV-4-11 cells modified for stable firefly Luciferase (ffLuc) expression^31^ or 5×10^4^ MOLM-13.ffLuc cells^32^ via tail vein on day 0. NK cell treatment was administered on D7 (10×10^6^cells) or D4, 7, and 10 (3×10^6^cells each). Mice were given D-Luciferin (3 mg) by intraperitoneal injection and bioluminescence (BL) measured using IVIS Spectrum (In Vivo Imaging System). Data were analyzed using Living Image Software (v 4.7.3; PerkinElmer, Waltham, MA). When indicated, peripheral blood was drawn via facial vein, red blood cells were lysed with eBioscience RBC Lysis Buffer (ThermoFisher) and the remaining cells analyzed with flow cytometry. Bone marrow and spleen were harvested and tissues analyzed with flow cytometry. Analysis of peripheral blood for cytokines (human (h) IL15, hTNFα, mouse (m) IL6, mIL1β) was performed with ELISA (R&D Systems). Mice were euthanized when they exhibited >20% weight loss, hind limb paralysis or moribund state as per protocol guidelines.

### Statistical analysis

All statistical analyses was performed using GraphPad Prism Software (v9.2.0). Our comparisons included more than three groups and ordinary one- or two-way analysis of variance (ANOVA) corrected using the method of Bonferroni. Data with variance of several logs of magnitude were log transformed (Y=log(Y)) before analyzing with ANOVA. Survival of mice was estimated by the Kaplan–Meier method and differences in survival between groups were calculated by the log-rank (Mantel–Cox) test.

## Results

### CD123-CARs are highly expressed on the NK cell surface

We considered NK cell biology in our design of 8 different NK-tailored CARs (Fig. 1A) to complement the common 4-1BB.ζ CAR.^33,34^ All CARs are comprised of an extracellular scFv targeting CD123.^25^ The hinge, transmembrane (TM), and intracellular portion of our CARs consisted of different combinations of activating co-receptors DAP10 and FcεRIγ, the co-stimulatory receptor 2B4, and the ζ chain of the T cell receptor (Fig. 1A). All CARs were expressed stably on the surface of primary human NK cells for at least two weeks in culture, with transduction efficiencies ranging from 21-98% (Fig. 1B). Representative flow cytometric plots are shown in Supplemental Fig. 1A. CARs encoding 2B4 or CD8α TM domains demonstrated higher transduction efficiencies (median(range), 89(53-98%) and 84(75-90%), respectively) than constructs containing FcεRIγ or DAP10 TM (62(21-77%) and 39(24-64%), respectively; Fig. 1B,C). 2B4 and CD8α TM domains also conferred optimal CAR surface density as estimated by comparative mean fluorescence intensities (MFI +/- SEM; 2B4-TM: 2158 +/- 242; CD8α-TM: 3254 +/- 970; FcεRIγ-TM: 827 +/- 151; DAP10-TM: 366 +/- 23; 2B4 and CD8α vs DAP10: p<0.0001; 2B4 vs FcεRIγ: p<0.01; CD8α vs FcεRIγ: p<0.0001; FcεRIγ vs DAP10: ns; Supplemental Fig. 1B). CAR-NK cells expanded 70-213 fold within 18 days of *ex-vivo* culture with no significant differences between generated CAR-NK cell populations (Supplemental Fig. 1C).

### CD123-CAR NK cells have antigen-specific anti-AML activity *in vitro*

We evaluated the target specificity of our CAR-NK cells using the CD123-positive MV-4-11 and CD123-negative Raji cell lines. When challenged with MV-4-11 in co-culture assays, all CAR-NK cells responded with enhanced cytokine secretion above that seen when using unmodified NK cells under identical conditions (mean percent (%) change of IFN-γ secretion; CAR-NKs(range): 65-313% vs unmodified: 14%, Supplemental Fig. 2). There was no difference in cytokine production of CAR(+) and CAR(-) NK cells after co-culture with CD123(-) targets (Supplemental Fig. 2). Next, we assessed CAR-NK cell cytotoxicity against CD123(+) target cells in 72-hour co-culture assays. We found that CARs with 2B4 or 4-1BB co-stimulatory and TCRζ signaling domains endowed NK cells with the greatest cytolytic activity against MV-4-11 at both 1:1 (mean % cytotoxicity +/- SEM; 2B4.ζ: 93.8 +/- 5%; 4-1BB.ζ: 89.9 +/- 6.8%) and 1:5 effector:target (E:T) ratios (82.9 +/- 11.6%; 93 +/- 2.6%, respectively; Fig. 1D). CD123(-) Raji cells were again used as controls. There was no difference between CAR-NK versus unmodified NK cell mediated cytotoxicity against Raji cells, confirming specificity (Fig. 1D).

### CD123-2B4.ζ CAR-NK cells have limited anti-AML efficacy *in vivo*

Given the superior anti-AML activity of 2B4.ζ and 4-1BB.ζ CAR-NK cells in our *in vitro* assays, we next evaluated their antitumor activity in a xenograft model of human AML. NSG mice were first engrafted with MV-4-11.ffLuc cells,^31^ then treated with CAR-NK or unmodified NK cells on day 7 (Fig 2A). Leukemic growth was measured with serial BLI. 2B4.ζ CAR-NK cells post injection had transient anti-AML activity, which translated into a significant survival advantage in comparison to other experimental groups (median survival (days); 2B4.ζ: 63 vs. 4-1BB.ζ: 56, p<0.01; 2B4.ζ vs. unmodified: 63 vs. 55, p<0.05) and untreated controls (2B4.ζ vs. no treatment: 63 vs. 58, p<0.01; Fig. 2B-D). Serial analysis of peripheral blood in our animals showed declining NK cell numbers in all evaluated mice irrespective of infused NK cell type (Fig. 2E). Planned bone marrow and spleen examination in a subset of mice demonstrated increasing percentages of leukemia from day 15 to day 22 in all treatment conditions, despite readily detectable NK cells (Supplemental Fig. 3). Consistent with BLI data, the percentage of leukemic cells was lower in both bone marrow and spleen samples of 2B4.ζ CAR-NK cell treated mice as compared to mice treated with UTD and 4-1BB.ζ CAR-NK cells (Supplemental Fig. 3).

**Figure 2.**
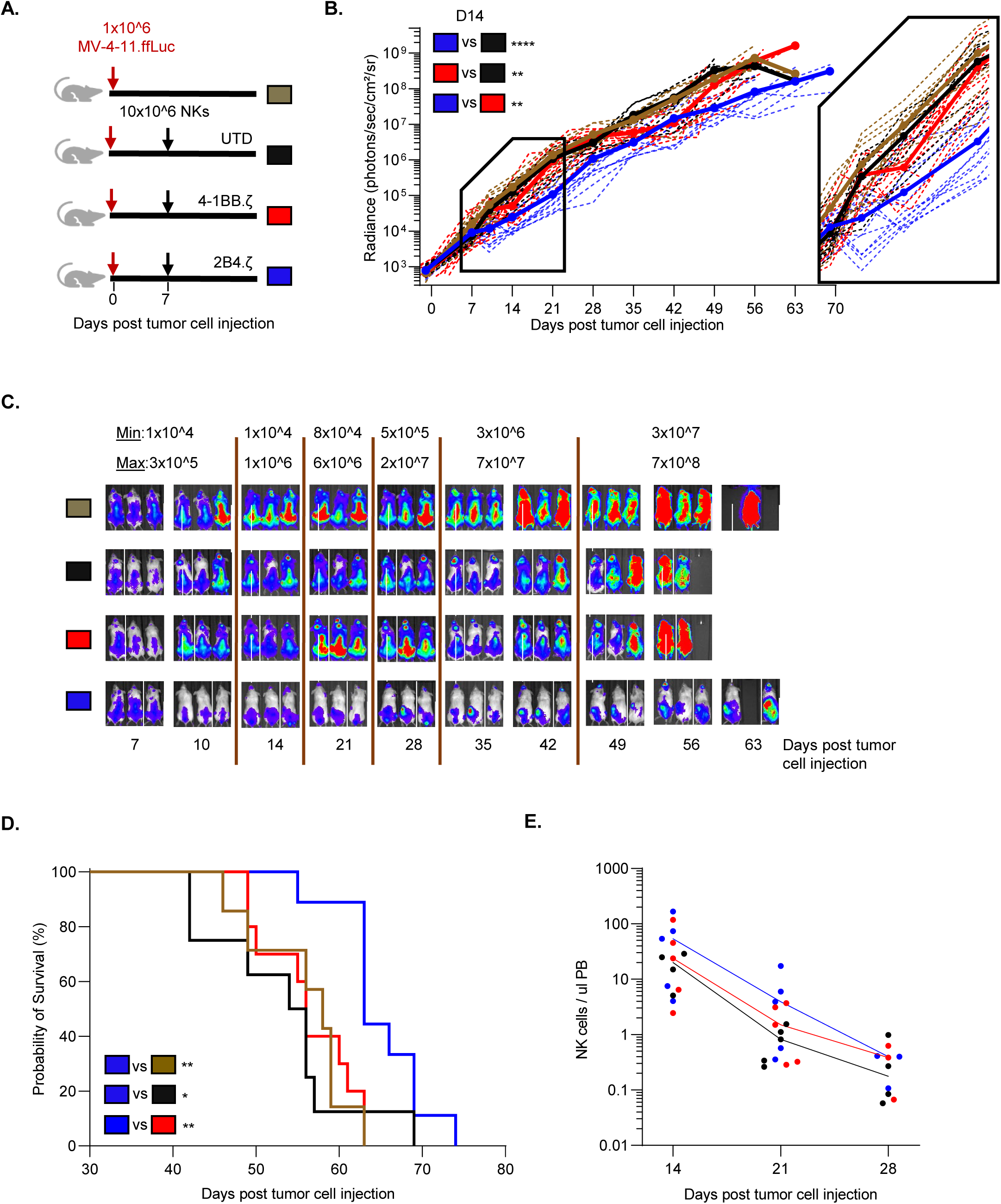
Anti-CD123.2B4.ζ CAR-NKs have transient anti-AML activity *in vivo*. (**A**) Schematic of MV-4-11 xenograft model. On day 0, NSG mice were injected via tail vein with 1×10^6^ CD123(+) MV-4-11 cells that express firefly Luciferase (MV-4-11.ffLuc cells). In treatment groups, 10×10^6^ NK cells were administered on day 7. Cohorts: untransduced/unmodified (UTD), 4-1BB.ζ CAR-NK, and 2B4.ζ CAR-NK. (**B**) Leukemia proliferation was monitored with bioluminescence imaging (BLI) and was recorded as photons/sec/cm²/sr; n=8-12 mice per group. Magnification of days 7-21 shown. (**C**) Representative images of 3 mice per condition. Minimum and maximum values of color scale are depicted at top [min-max]. (**D**) Kaplan–Meier survival analysis of MV-4-11 xenografts (n=8-12 mice per condition). (**E**) Mouse peripheral blood (PB) collected at indicated time points and analyzed via flow cytometry. Each dot represents a single mouse. Solid line: median. At later time points, NK cell count was undetectable for all groups and is not plotted (*p<0.05; **p<0.01; ***p<0.001).

### Armoring NK cells with secreted IL15 enhances anti-AML functionality *in vitro*

Having demonstrated that 2B4.ζ CAR-NK cells have anti-AML cytolytic capacity with short persistence, we next explored if transgenic expression of IL15 in CAR-NK cells would support sustained anti-AML activity. To accomplish this, we cloned a sequence encoding human IL15 downstream of an IRES element into our 2B4.ζ CAR vector. We simultaneously generated a second retroviral vector encoding IL15 and the fluorescent molecule mOrange as a control with IL15, but not CAR expression (Fig. 3A). We verified transduction by measuring vector copy number (VCN) per cell (median [range]; 2B4.ζ: 10[5.8-14]; 2B4.ζ/secretory(s)IL15: 4.1[3.8-4.7]; sIL15/mOrange: 16.6[4.8-19.2]; Fig. 3B). We found no significant difference in VCN/cell or in CAR expression between NK cells transduced with 2B4.ζ or 2B4.ζ/sIL15 encoding retroviral vectors (range CAR(+); 2B4.ζ: 70-97% vs. 2B4.ζ/sIL15: 81-96%, p>0.99; MFI +/- SEM: 2,566 +/- 531 vs. 2,089 +/- 424, p>0.99; Fig. 3C, Supplemental Fig. 4A). Expression of IL15 was measured with qRT-PCR (Supplemental Fig. 4B) and IL15 secretion was confirmed by ELISA (Fig. 3D).

**Figure 3.**
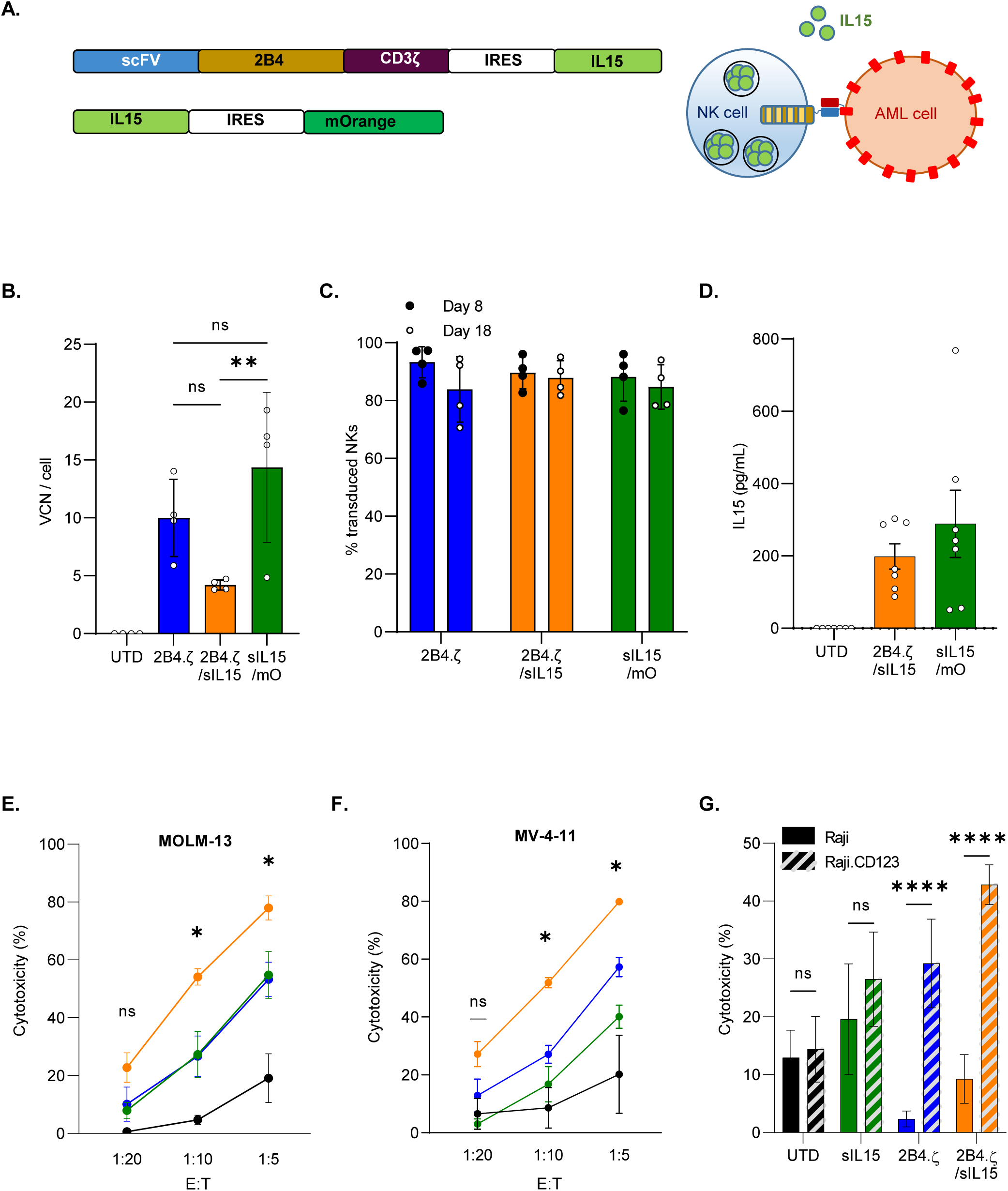
Simultaneous 2B4.ζ CAR expression and IL15 secretion strengthens NK cell cytotoxicity. (**A**) Schema of vectors and IL15 secretion from CAR-NK cells. (**B**) Quantification of retroviral vector copy number (VCN) in transduced NK cells. UTD NK cells served as negative controls (n=4 donors). (**C**) Percentage (%) of transduced NK cells in cultures on D8 (black circle) and D18 (white circle, n=4 donors). (**D**) NK cell supernatant was used for quantification of IL15 by ELISA (n=7 biological replicates using 4 donors). BL based cytotoxicity assays performed using the CD123(+) AML cell lines MV-4-11(**E**) and MOLM-13 (**F**) with ffLuc expression (n=4 donors). Asterisks indicate 2B4.ζ/sIL15 vs 2B4.ζ comparison. (**G**) Bar graph comparing percent (%) cytotoxicity of NK cells against Raji (CD123(-): solid), and Raji.CD123 (CD123(+): diagonal stripe) cancer cell lines at 1:10 E:T ratio (n=4 donors). For panels B-G, mean +/- SEM represented (*p<0.05; **p<0.01; ***p<0.001, ****p<0.001).

We evaluated short-term cytotoxicity of 2B4ζ/sIL15 CAR-NKs against the CD123(+) MV-4-11, MOLM-13, and Raji.CD123 cell lines. The parental CD123(-) Raji line was used as a negative control. When compared to 2B4.ζ, the 2B4ζ/sIL15 CAR-NKs had higher cytotoxicity against CD123+ targets (Fig. 3E-G). Both 2B4.ζ and 2B4.ζ/sIL15 demonstrated antigen-specific cytotoxicity as Raji.CD123 were more effectively killed than parental (CD123 negative) Raji cells (Fig. 3G). Target cell CD123 and IL15Rα expression were quantified in order to evaluate for any effect of CD123 surface density or IL15 *trans* presentation on NK cell cytotoxicity (Supplemental Fig. 5A). Differences in measured CD123 surface density did not correlate with observed short-term cytotoxicity, underlining the existence of additional complex mechanisms affecting NK cell activation. Similarly, differences observed in IL15Rα expression did not correlate with cytotoxicity (Supplemental Fig. 5B).

### Transgenic expression of IL15 potentiates the activation, persistence, and long-term cytolytic activity of CAR-NK cells

Using a model of chronic antigen stimulation (Fig. 4A), we evaluated the immune phenotype of our CAR-NK cells at baseline, after 12 hours, and on day 10 (D10) of co-culture with MV-4-11. Cells were counted daily, and AML repleted to maintain a 1:1 E:T ratio. We used flow cytometry to measure surface expression of markers in two different panels (panels A and B), then performed hierarchical clustering of NK cell subsets (Fig. 4B,C, Supplemental Fig. 6). In panel A, we observed similar population distributions in the 2B4.ζ CAR and unmodified NK cells with cluster shifts from earlier (12h) to later (D10) time points. Specifically on D10, there was an increase in the percentage of 2B4.ζ and unmodified NK cells populating clusters defined by lower surface expression of activating receptors NKG2D and NKp30 (clusters *4, 5, 23*; Fig 4B,C, Supplemental Fig. 7). For 2B4.ζ/sIL15 CAR-NK cells, only minimal changes in population density of these subsets were observed on D10 (Fig 4B,C). The unique 2B4.ζ/sIL15 immunophenotype is highlighted by comparison of the MDS (global) and UMAP (cluster-specific) plots in Supplemental Fig 8.

**Figure 4.**
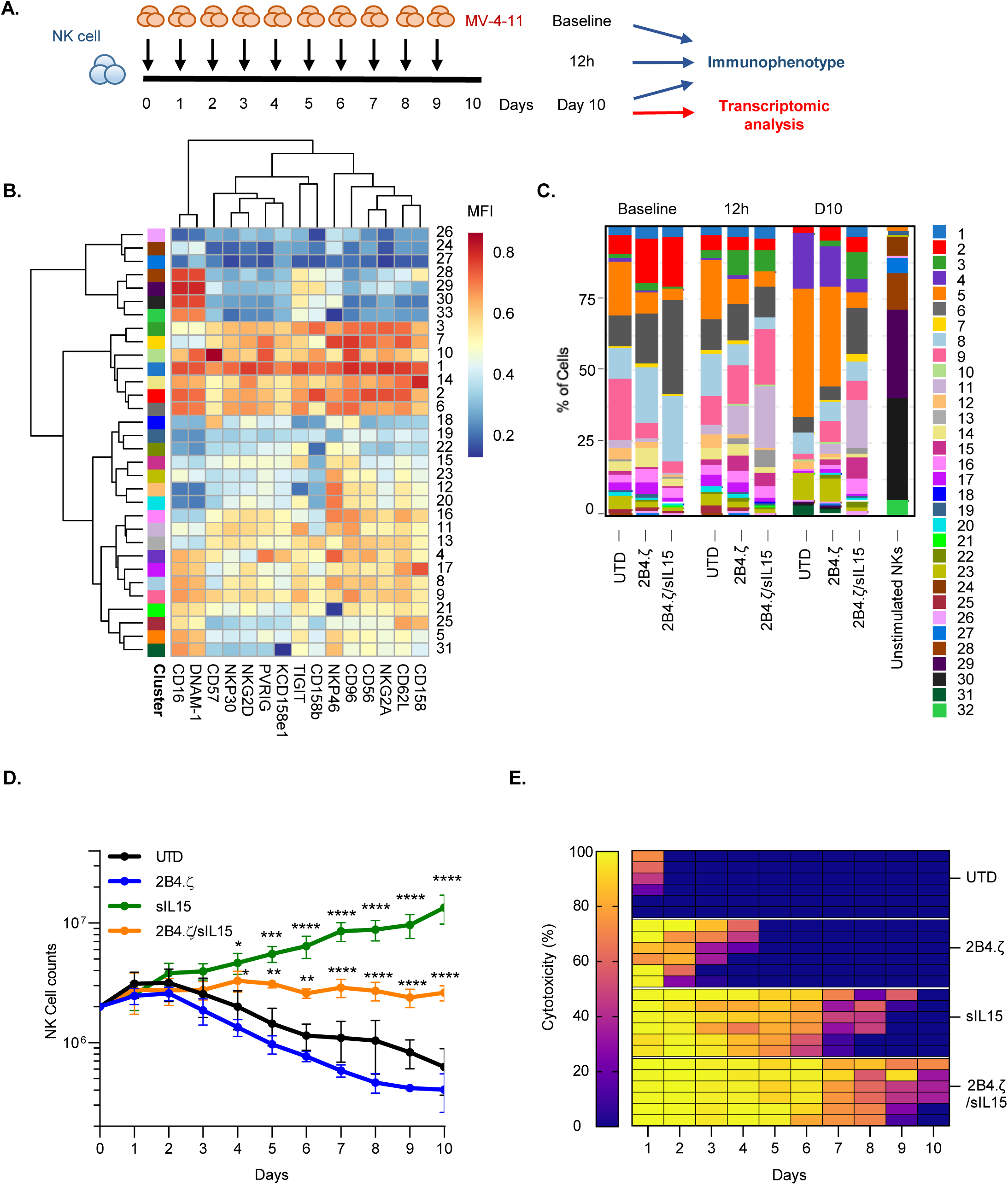
IL15 maintains NK cell activated phenotype in a model of chronic antigen stimulation. (**A**) Schematic representation of our serial stimulation assay. Day 0 was the day of the initial seeding of the co-culture; day 1 the first and day 10 (D10) the last day of cell quantification. Immunophenotypic analysis of effector and target cells performed at baseline (before co-culture), 12 hours after 1^st^ stimulation, and on D10 (n=1 donor). Transcriptomic analysis performed on D10 (n=3 donors). (**B**) Heatmap of flow cytometry data showing expression of 15 different NK cell surface markers. Heatmap coloring represents arcsinh transformed median marker intensity. (**C**) Bar plots of relative abundance of the 32 population subsets found in each sample. (**D**) NK cell counts over a period of 10 days. Initial seeding count was 2 million NK cells (mean +/- SEM; n=3 donors). Asterisks indicate 2B4.ζ/sIL15 vs 2B4.ζ and sIL15 vs UTD comparison. (**E**) Heat map of the percent (%) NK cell cytotoxicity. Each column represents the specific day and each row a unique biological replicate (n=3 donors).

Clustering of NK cells based on expression of markers included in our second receptor panel (panel B) again supported the maintained NK cell activation to D10 in 2B4.ζ/sIL15 CAR-NK cells (predominant clusters *9, 15, 17* and minor clusters *1, 4, 11, 19, 22, 24*; Supplemental Fig. 6,9). Differences were again observed in 2B4.ζ and unmodified NK cells on D10 compared to earlier time points (increasing percentage of cells populating clusters *16, 18, 20, 23* and decreasing percentages of *4, 9, 15, 17*; Supplemental Fig. 6). Clusters *4*, *9, 15, and 17* expressed higher levels of LFA-1, CD69, TRAIL, TIM-3, NKG2A, and KLRG1 compared to clusters *16*, *18*, *20*, and *23* (Supplemental Fig. 9). Negligible differences were observed in PD-1, LAG3, FASL, 2B4, and NKG2C expression. Taken together, 2B4ζ/sIL15 CAR-NKs had a higher percentage of NK cells populating clusters defined by higher surface expression of LFA-1(adhesion/activation receptor), CD69 (activation marker), TRAIL (death receptor), and Tim-3 (commonly upregulated after NK cell activation), as well as NKG2A and KLRG1 (inhibitory receptors; Fig. 4B,C, Supplemental Fig. 6-10). Overall, the data from both antibody panels suggests that with continuous antigen stimulation, IL15 preserves an activated CAR-NK cell phenotype associated with high cytotoxicity. We also analyzed the expression of NK cell activating and inhibitory ligands. AML cells expressed high levels of MICA/MICB (NKG2D ligands), CD112 and CD155 (DNAM1, PVRIG, CD96, and TIGIT ligands) and PDL1 (PD-1 ligand), but not ULBP1 (NKG2D ligand) or galectin-9 (Tim-3 ligand; Supplemental Fig. 11).

Next, we evaluated NK cell cytotoxicity and persistence in our model of chronic antigenic stimulation (Fig. 4A). While unmodified and 2B4.ζ CAR-NK cells sharply diminished in number over the course of this assay, NK cells genetically engineered to express IL15 (2B4.ζ/sIL15 and sIL15) survived throughout the experiment (n=3; Fig. 4D). As predicted by the short-term cytotoxicity assay, 2B4.ζ CAR-NK cells had high initial anti-AML activity. However, their killing potential decreased as 2B4.ζ CAR-NK cell counts declined. Cytotoxicity was completely abrogated by day 5 (Fig. 4E). IL15 activation alone (without CAR expression) also had an early anti-tumor effect that was not sustained. Only the combination of CAR expression and IL15 secretion led to continued NK cell mediated anti-AML cytotoxicity (Fig. 4E, Supplemental Fig. 12).

### NK cells secreting IL15 exhibit a highly proliferative and activated transcriptomic signature after chronic antigen stimulation

We evaluated the transcriptional programs of our NK cells on the tenth day of continuous antigen stimulation. NK cells were isolated, and RNA libraries prepared and used for RNAseq analysis. Samples clustered by IL15 secretion, with overlap between 2B4.ζ/sIL15 and sIL15 conditions. These clearly separated from NK cells that did not secrete IL15 (UTD and 2B4.ζ; Fig. 5A). Differential gene expression analysis (DESeq2) of 2B4.ζ/sIL15 vs 2B4.ζ and sIL15 vs UTD NKs identified in the IL15-secreting NK cells differences in expression of genes in the pathways of DNA replication, cell cycle progression, and NK cell mediated cytotoxicity (Fig. 5B,C). KEGG enrichment analysis revealed cell cycle progression as the top-ranked pathway for both comparisons (Fig. 5C). No biologically relevant pathways were found to be significantly enriched when comparing IL15-secreting CAR-NK cells to non-CAR sIL15 NKs (Supplemental Fig. 13A, B). Hierarchical clustering of differentially expressed genes also revealed upregulation of genes involved predominantly in cell cycle progression, chemokine, and cytokine signaling in IL15-secreting NK conditions (Fig. 5D, Supplemental Table 2). We used this dataset to evaluate differential expression of molecules of biological relevance. 2B4.ζ/sIL15 and sIL15 NK cells had higher expression of genes encoding for NK activating receptors (*NCR2*, *NCR3*, *KLRC2*, *KLRC4-KLRK1*, *CD226, FCGR3A*), adaptor molecules (*FCER1G*), death receptor ligands (*TNFSF10*), granzyme (*GZMA*), proinflammatory cytokines and chemokines (*IFNG*, *CCL1, CCL3*, *CCL4*, *XCL1*, *XCL2, CCL3L3*), activation markers (*CD69*), proliferation markers (*MKI67*), anti-apoptosis regulators (*BCL2*), and adhesion molecules (*ITGB2, CD2, CD53*; Fig. 5E, Supplemental Fig. 13C). We also observed upregulation of select inhibitory receptors (*NKG2A*, *CEACAM1*, *LILRB1*) including inhibitory Killer Ig-Like Receptors (KIRs; *KIR2DL1, KIR2DL2, KIR2DL3, KIR2DL4, KIR3DL1*) as well as “checkpoint” molecules (*HAVCR2*, *TIGIT*; Fig. 5E, Supplemental Fig. 13C). KIR are subject to extensive regulatory splicing and there is an inability to distinguish KIR with identical extracellular domains but functionally disparate intracellular tails by flow cytometry, making transcriptional analysis necessary and complementary for study of KIR expression. Analysis of the chemokine receptor expression showed upregulation of *CCRL2, CCR1, CCR5 and CCR6* with similar or lower expression of *CX3CR1, CXCR3* and *CXCR4* in IL15-secreting NKs (Supplemental Fig. 13C).

**Figure 5.**
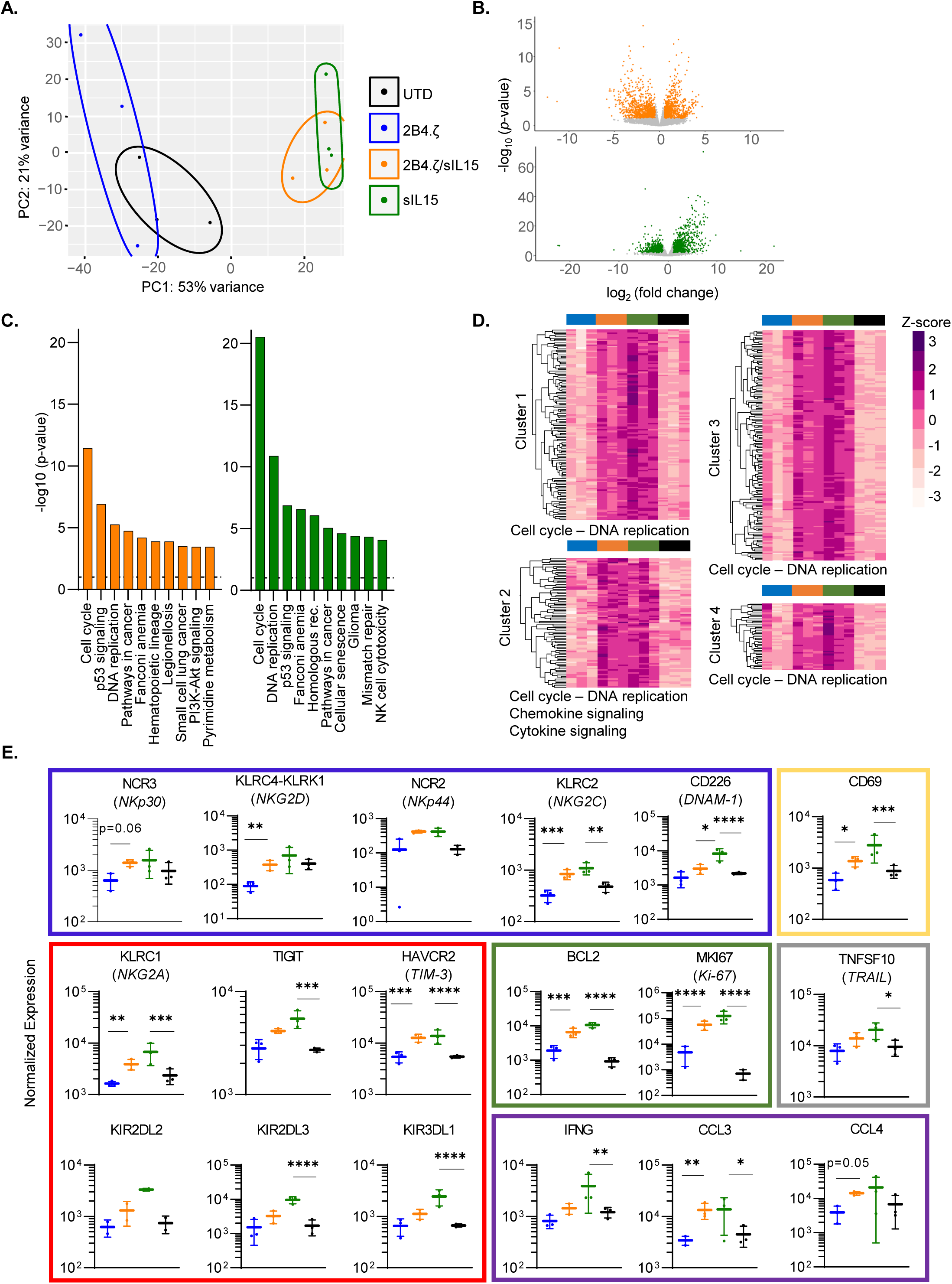
IL15 stimulation of NK cells upregulates genes important to cell cycle progression, NK cell activation, and cytotoxicity. (**A**) Principal component analysis (PCA) of NK cell transcriptome on D10 of the chronic antigen stimulation. Each dot represents a unique NK cell donor (n=3 donors). (**B**) Volcano plots representing differentially regulated genes in 2B4.ζ/sIL15 compared to 2B4.ζ (orange) and sIL15 compared to unmodified cells (green). Grey dots are those not meeting criteria: p-value ≥ 0.05; fold change > 2 or <1/2. (**C**) Bar plot depicting the top 10 significantly different KEGG 2021 Human gene Set Enrichment pathways Dotted line at p-value of 0.05. (**D**) Heatmap and hierarchical clustering performed on differentially expressed genes. Biologically relevant clusters and enriched pathways (shown on the bottom of the cluster) represented. Z-score scale bar at right. (**E**) Dot plots displaying the normalized expression of genes encoding activating receptors (blue rectangle), inhibitory receptors/checkpoint molecules (red), proliferation/anti-apoptosis markers (green), inflammatory mediators (purple), death receptor ligands (grey) and activation (yellow) markers (n=3 independent donors; *p<0.05; **p<0.01; ***p<0.001; ****p<0.0001).

### Constitutive IL15 expression improves NK cell *in vivo* persistence, but causes lethal toxicity

We next evaluated sIL15 NK cells in two AML xenograft models. In our first experiment, we used our previous NK cell dosing regimen. Surprisingly, MV-4-11 engrafted mice treated with 10×10^6^ IL15-secreting NK cells had early mortality (median survival(days); *2B4.ζ/sIL15*: 25 ; *sIL15*: 21; n=5 mice each group; Supplemental Fig 14). Therefore, we subsequently compared a single decreased dose of 2B4.ζ/sIL15 CAR-NK cells with multiple doses of 2B4.ζ CAR-NK cells (Fig. 6A). 2B4.ζ/sIL15 CAR-NK given as a single injection of 3×10^6^ cells and three doses of 3×10^6^ 2B4.ζ CAR-NK cells maintained transient AML control (Fig. 6B,C). Three doses of 2B4.ζ CAR-NKs prolonged survival, as compared to mice in all other treatment groups (median survival(days); 2B4.ζ: 71 vs UTD: 48 and untreated: 49, p<0.001; Fig. 6D). In contrast, even at the lower dose, toxicity and premature death of mice treated with IL15-secreting NK cells again occurred (median survival (days); 2B4.ζ/sIL15: 26 vs untreated: 49, p<0.001; sIL15: 21 vs untreated: 49, p<0.001; Fig. 6D). Analysis of peripheral blood showed *in vivo* expansion of 2B4.ζ/sIL15 and sIL15 NKs, with declining NK cell counts again observed in non-IL15 secreting NK cell cohorts (Supplemental Fig 15A). Bone marrow and spleen analysis at necropsy of mice treated with 2B4.ζ/sIL15 CAR-NKs revealed high numbers of infiltrating NK cells (Supplemental Fig. 15B,C). We identified increasing systemic hIL15 in mice treated with 2B4.ζ/sIL15 and sIL15 NKs (Supplemental Fig. 15D). In addition, high levels of hTNFα, low levels of mIL1β and negligible levels of mIL6 were detected in the blood of mice treated with 2B4.ζ/sIL15 CAR-NK cells (Supplemental Fig. 15E-G).

**Figure 6.**
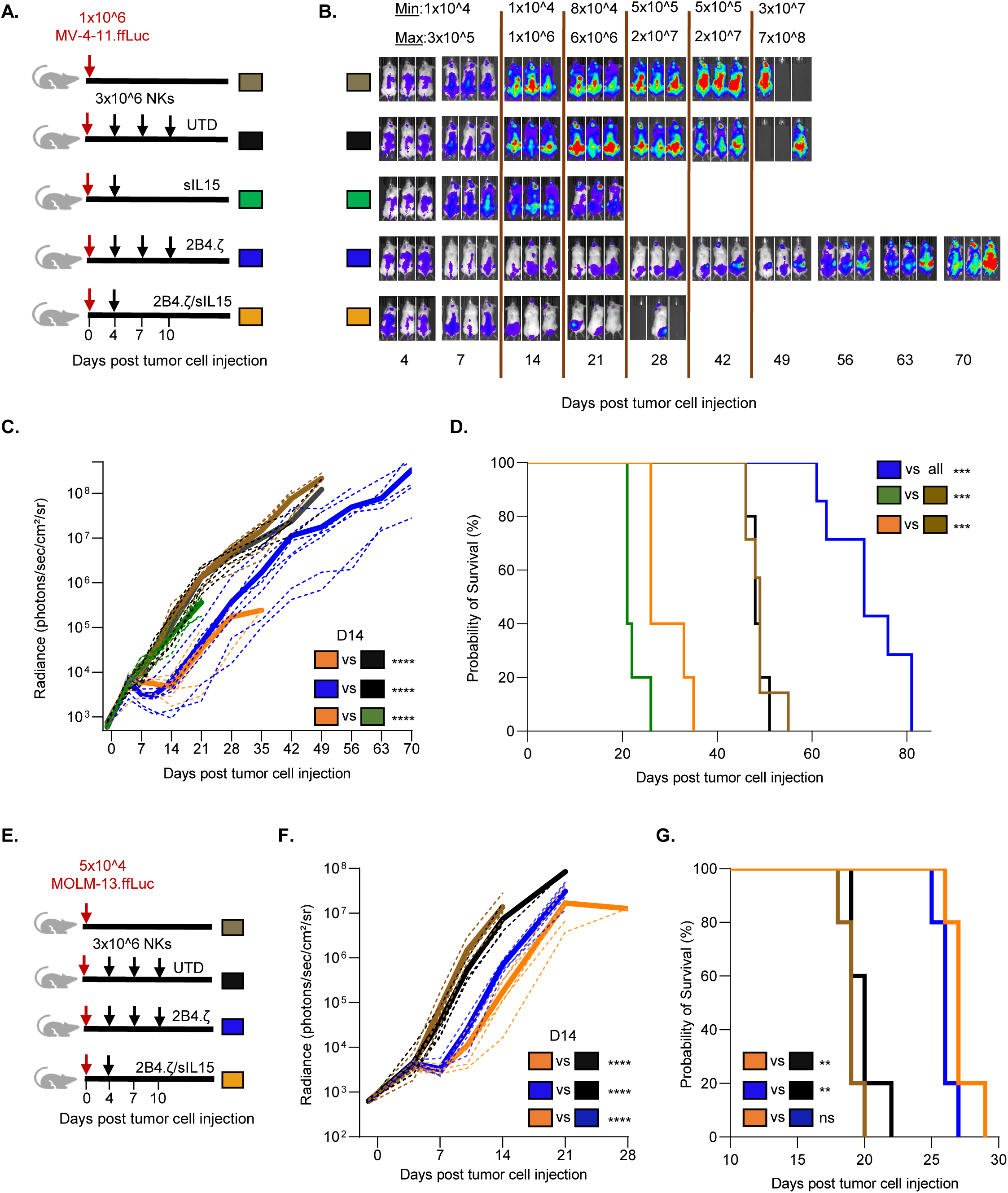
IL15 secreting CAR-NK cells can cause lethal toxicity in an AML xenograft model. (**A**) Schematic of NK cell dosing in MV-4-11.ffLuc model. (**B**) MV-4-11 proliferation was monitored with BLI. Representative images of mice. The minimum and maximum values of the color scale are indicated.[min-max] (**C**) Leukemia proliferation was monitored with bioluminescence imaging (BLI) and was recorded as photons/sec/cm²/sr . Dotted lines: individual mice, Solid lines: Mean (n=5-7 mice per group). (**D**) Kaplan–Meier survival analysis. (**E**) Schematic of NK cell dosing in MOLM-13.ffLuc model. (**F**) MOLM-13 proliferation was monitored with BLI (n=5 mice per group). (**G**) Kaplan–Meier survival analysis. (*p<0.05; **p<0.01; ***p<0.001; ****p<0.0001).

We further evaluated our CAR-NK cells in a more aggressive MOLM-13 xenograft model (Fig. 6E). A single dose of 3×10^6^ 2B4.ζ/sIL15 CAR-NK cells and three doses of 3×10^6^ 2B4.ζ CAR-NK cells had equivalent antitumor control (Fig. 6F, Supplemental Fig. 16A). In contrast to the lethal toxicity seen in the MV-4-11 model, both treatment strategies prolonged survival in MOLM-13 engrafted mice (median survival (days); 2B4.ζ: 26 vs UTD: 20, p<0.01; 2B4.ζ/sIL15: 27 vs UTD: 20, p<0.01; Fig. 6G). Analysis of peripheral blood at necropsy showed circulating systemic hIL15 and hTNFα, but negligible levels of mIL1β and mIL6 (Supplemental Fig. 16B-E).

## Discussion

Herein we describe the generation and functional evaluation of chimeric receptors targeting the AML-associated antigen CD123 and expressed in primary human NK cells. The molecular domains distal to the extracellular scFv consisted of various combinations of NK-specific activation moieties. We identified a 2B4.ζ CAR as having similar *in vitro* functionality to the well-studied 4-1BB.ζ CAR,^35^ with high surface expression, antigen-specific activation, and cytotoxicity.

The *in vivo* anti-tumor effect was more pronounced when AML-engrafted mice were treated with 2B4.ζ CAR-NK cells as compared to those treated with 4-1BB.ζ CAR-NK cells, which suggests a potential additive effect of 2B4 and CD3ζ signaling. However, this *in vivo* effect was short-lived and was accompanied by circulating NK-cell decline. We thus engineered our 2B4.ζ CAR-NK cells with an IL15 transgene to promote IL15-mediated activation, proliferation, and survival. Secretion of IL15 stimulated CAR-NK cell expansion both *in vitro* and *in vivo*, and enhanced short- and long-term anti-AML cytotoxicity. This bolstered activation profile was confirmed by immunophenotypic and transcriptomic analysis in the setting of chronic stimulation. However, in one *in vivo* AML model, constitutive IL15 secretion caused dramatic NK-cell expansion, high levels of circulating human pro-inflammatory cytokines, and was associated with early death. Treatment with three doses of non-IL15-secreting CAR-NK cells prolonged survival without systemic toxicity, but anti-tumor efficacy was transient.

CARs are used to enhance and redirect immune effector cells against cancer cells. CAR-T cells have been extensively investigated in preclinical and clinical models of AML.^9^ However, there are limited preclinical animal studies^36,37^ and only 3 active clinical trials (NCT04623944, NCT05008575, NCT02742727) testing CAR-NK cells as AML therapy. There is a single completed anti-AML CAR-NK cell trial with evaluable data (NCT02944162). This study tested an engineered anti-CD33 CAR-NK92 cell line in 3 patients with relapsed and refractory AML. The CD33-CAR NK92 cell infusion was safe, but the treatment had minimal antitumor efficacy.^38^ We aimed to enhance the molecular functionality of our CAR by using NK cell-specific receptor domains. We identified the 2B4TM-2B4-CD3ζ CAR as optimal for NK cell expression and activation. The end-result of 2B4 downstream signaling includes synergy with other NK activating receptors.^17,39^ Protein interactions including 2B4 occur inside membrane (“lipid”) rafts. We chose to use the 2B4 transmembrane domain to localize the CAR within lipid rafts,^40^ and found that CARs containing a 2B4 transmembrane domain had high surface density and stable CAR expression. Tandem CD3ζ was also important, highlighted by the observed functional differences between 2B4.ζ and 2B4 CAR-NK cells.

To our knowledge, we describe the first study investigating both the *in vitro* and *in vivo* anti-AML functionality of peripheral blood derived CAR-NK cells (PB-NKs) targeting CD123. We chose the peripheral blood (PB) of healthy donors as the source of our CAR-NK cells. PB-derived NK cells (PB-NKs) can be isolated through apheresis and expanded in large scale using feeder cells.^26^ PB-NKs are functionally mature, with high activating receptor expression and cytotoxic potency. PB-NKs also display higher levels of Killer Ig-Like Receptors (KIRs) compared to other NK cell products derived from alternative sources. KIR expression is indicative of more complete NK cell licensing.^41^ Donor-derived, allogeneic PB-NK cells have therapeutic potential due to their relative safety, immediate availability once manufactured and stored, and reduced manufacturing costs as compared to per-patient manufacture of autologous cell therapy products.^12,13^ Historically, one challenge facing PB-NK cell engineering was that of poor viral and non-viral genetic modification.^42^ With our method, we are able to achieve high levels of PB-NK transduction. All of our CARs were stably expressed on the surface of primary NK cells, though inter-CAR variability in surface density was observed.

NK cells have a natural lifespan of approximately 2 weeks in humans. The success of adoptively transferred cellular therapies for cancer is determined, in part, by effector cell persistence. One strategy to support *in vivo* CAR-NK cell survival is to engineer constitutive expression of activating cytokine.^43^ This approach has been shown to be safe in a clinical trial using CD19-CAR NK cells against CD19+ lymphoid malignancies.^44^ This demonstrated safety profile motivated us to also test transgenic IL15 expression with a goal of enhanced *in vivo* CAR-NK cell persistence. In our study, treatment with IL15 secreting CAR-NK cells caused early death in mice engrafted with MV-4-11 AML. A likely cause of the observed systemic toxicity is severe inflammation due to the dramatic NK cell proliferation associated with high levels of circulating IL15 and other proinflammatory cytokines. A CRS-like syndrome triggered by murine monocytes or other immune cells is unlikely due to our inability to detect common murine proinflammatory cytokines. IL15 stimulation of accelerated leukemic growth was not observed. Our observation of unexpected toxicity begs for expanded study of novel ways to stimulate CAR-NK cell cytokine receptor pathways while circumventing systemic cytokine delivery. Localization of IL15 with membrane tethering^45^ or targeted delivery through the use of oncolytic viruses^46^ are alternative therapeutic strategies. Controllable cytokine expression using engineered inducible systems and safety switches also holds promise. Specific activation of intrinsic gamma-cytokine receptor signaling without the use of a pharmacologic agent is another strategy that has the potential to sustain NK cell function with an improved safety profile.

In conclusion, our data underscore the critical nature of IL15 signaling as a stimulant of CAR-NK cell long-term survival and anti-AML functionality. Our *in vivo* data highlight the limitations in murine modeling and the necessity for caution if the use of IL15 secreting NK cells is planned in human trials of CAR-NK cell adoptive transfer. Continued therapeutic development is imperative, to include testing of alternate and potentially safer methods of cytokine pathway activation.

## Author’s contributions

IC and CLB conceived of the work and developed the methodology. IC, AM, JR, AT, RR, AF, CR, SY, EES, and CLB performed the experiments. IC, WJH, MK, and CLB analyzed the data. IC, RV, and CLB performed statistical analysis. IC and CLB interpreted the data and wrote the manuscript. All authors revised the manuscript.

## Acknowledgements

We would like to thank Dr. Stephen Gottschalk and Dr. Peter Chockley for helpful discussion, review, and advice. We also thank the Johns Hopkins University Genetic Resources Core Facility for DNA sequencing, the Experimental and Computational Genomics Core for RNAseq data generation, and the Bloomberg-Kimmel Institute Cytometry Center for multiparameter flow cytometry data acquisition.

## List of Abbreviations

AML: Acute Myelogenous Leukemia
BL: Bioluminescence
CAR: Chimeric Antigen Receptor
CRS: Cytokine Release Syndrome
D10: day 10
E:T: effector-to-target
ffLuc: firefly Luciferase
GVHD: graft-versus-host disease
h: human
HCT: Hematopoietic Cell transplantation
H: hinge
IC: intracellular
ICANS: Immune Effector Cell Associated Neurotoxicity Syndrome
IL: Interleukin
KIR: Killer Ig-Like Receptor
m: mouse
MFI: mean fluorescence intensity
PB: peripheral blood
PB-NKs: peripheral blood derived NK cells
PBMC: peripheral blood mononuclear cells
s: secretory
SEM: standard error of mean
scFV: single chain variable fragment
TM: transmembrane
UTD: untransduced/unmodified
VCN: vector copy number

---

**Supplemental Figure 1.**
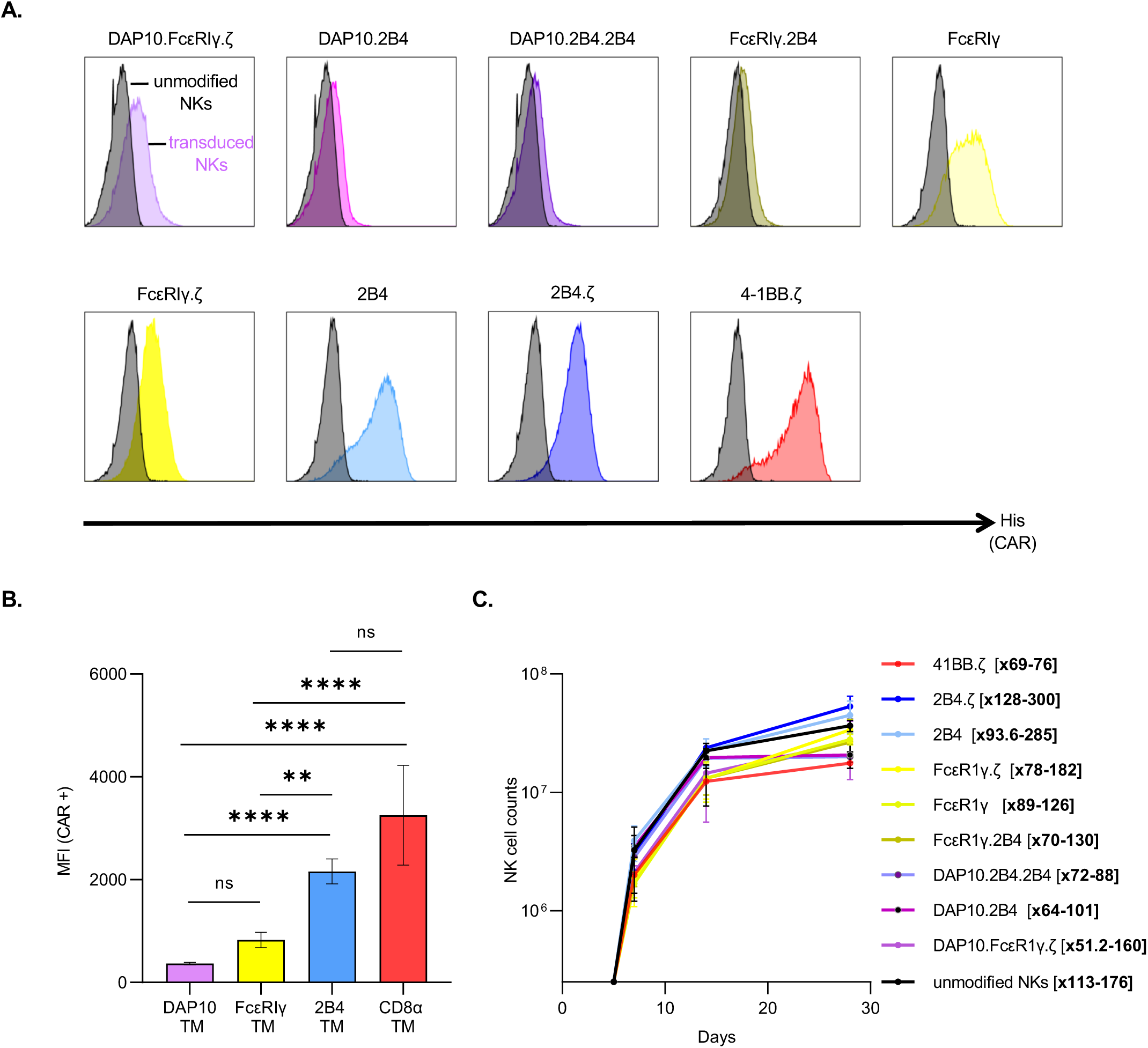
CAR expression and proliferative rate of CAR-NK cells. (**A**) FACS plots representing the gating strategy used to identify CAR+ NK cells. (**B**) Bar plot comparing the mean fluorescence intensities +/- SEM of CARs with indicated transmembrane (TM) domains. (**C**) Rate of expansion of our CAR-NK cells calculated with manual counting of absolute number. Initial seeding count was 250,000 cells. Fold expansion indicated in [ ] to the right of legend. Mean +/- SEM (n=3 donors).

**Supplemental Figure 2.**
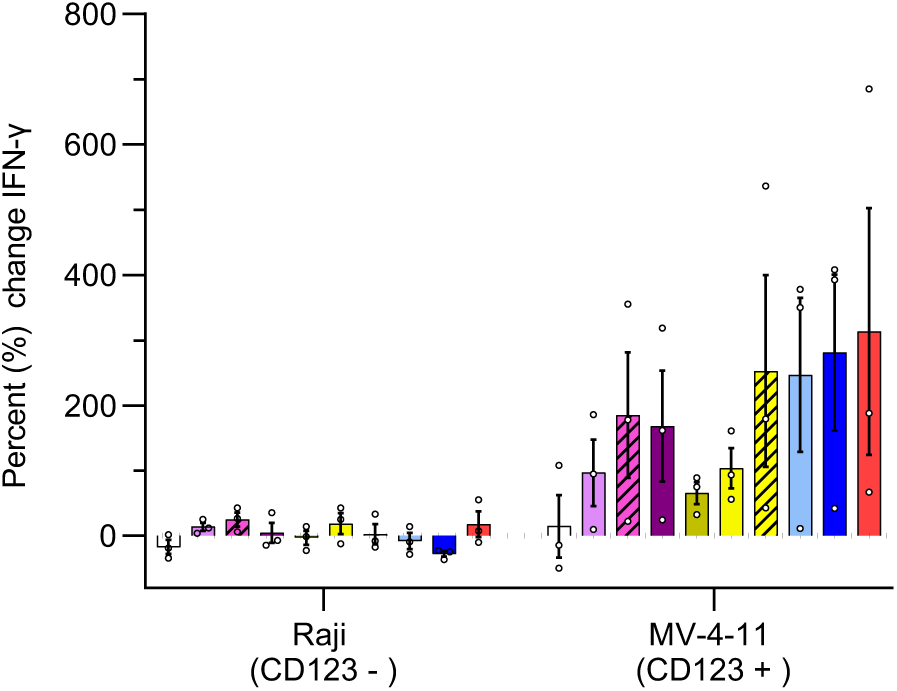
Anti-CD123 CAR-NK cells are activated by CD123+ cells. Percent (%) change of IFNγ secretion from baseline measured at 24-hours in co-culture assays of indicated CAR-NKs with Raji [CD123(-)] and MV-4-11 [CD123 (+)] cancer cell lines. IFNγ secretion was measured with ELISA. Each bar representative of the mean plus or minus the standard error of mean (+/- SEM); each dot is representative of an individual NK cell donor.

**Supplemental Figure 3.**
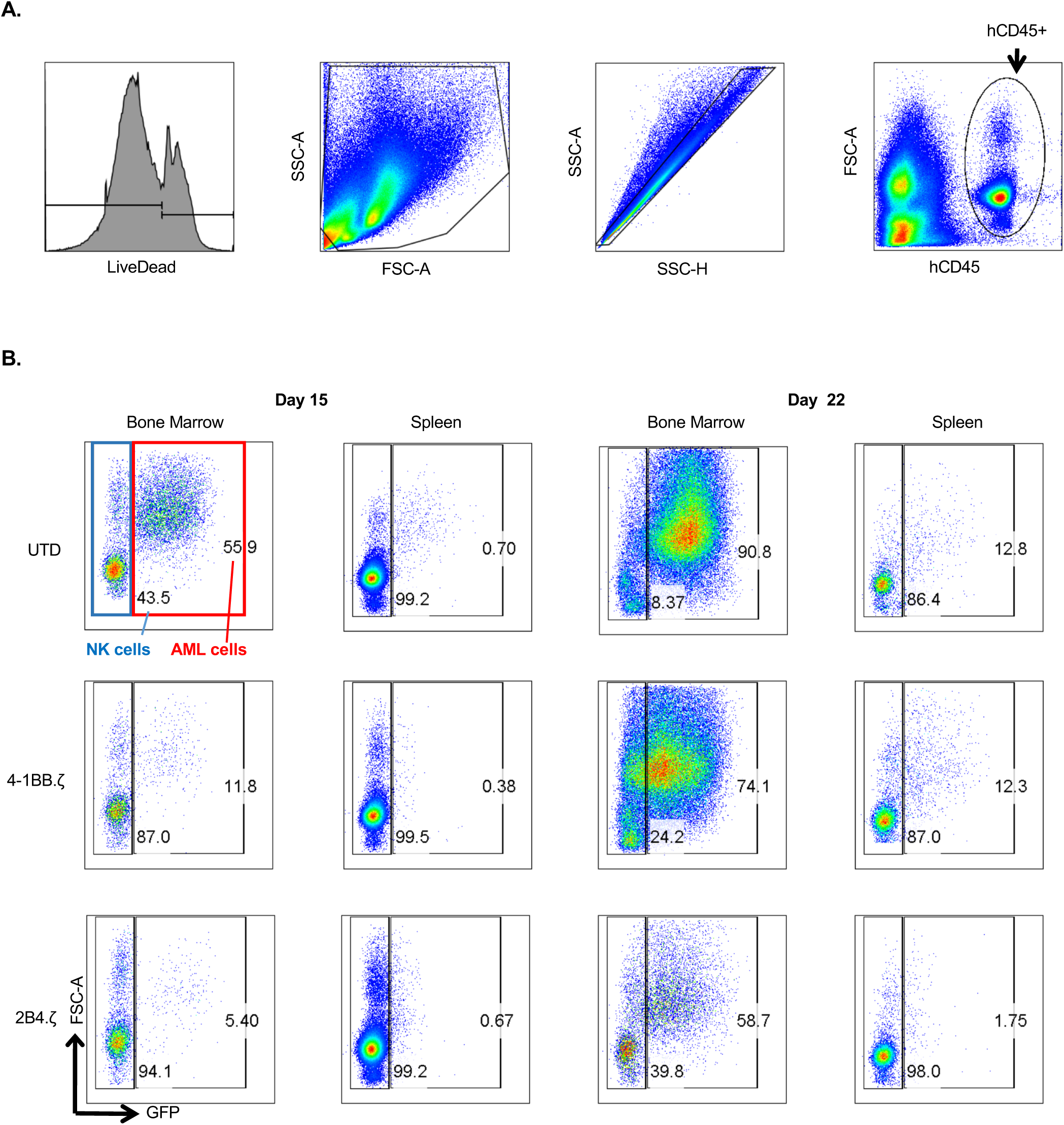
NK and leukemia cell percentages in the bone marrow and spleen of MV-4-11 engrafted mice on experimental days 15 and 22. **A.** Gating strategy used to identify hCD45+ cells in all *in vivo* experiments. LiveDead negative cells (alive) → excluding cellular debris → Single cells→ human CD45 positive (+) cells (human cells). **B.** FACS plots of hCD45(+) cells in the bone marrow and spleen of mice on day 15 (8 days after NK cell injection) and on day 22 (15 days after NK cell injection). MV-4-11.ffLuc cells (red rectangle) are GFP positive. Numbers represent percentages of NK or AML cells. One mouse analyzed per condition per day.

**Supplemental Figure 4.**
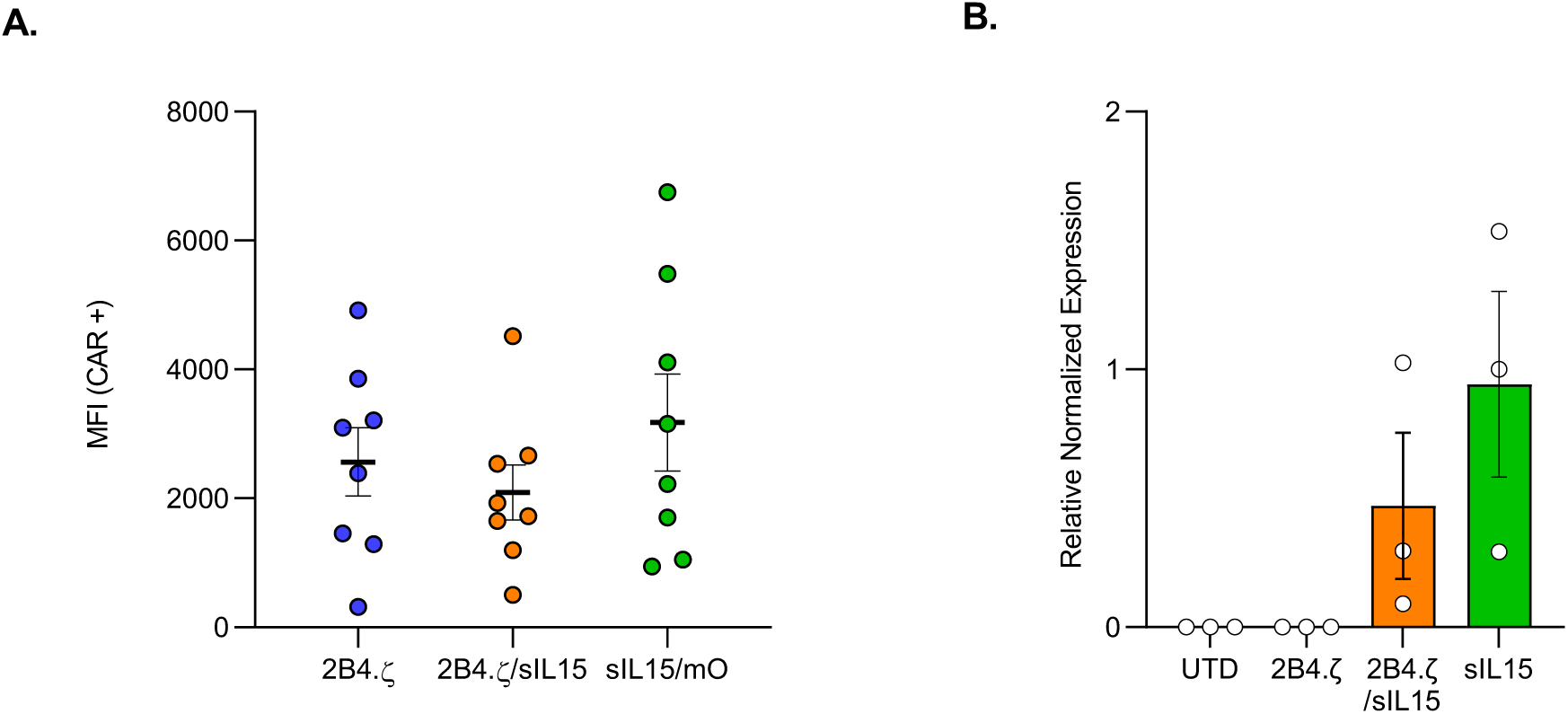
Expression of CARs and IL15 in NK cells. (**A**) Mean fluorescence intensity (MFI) of CAR (2B4.ζ and 2B4.ζ/sIL15) or mOrange (sIL15/mO) expression on transduced NK cells (n=4 donors measured at 2 time points). (**B**) Transgenic IL15 expression measured with Real-Time quantitative PCR. Normalized gene expression plotted relative to donor #3 sIL15 condition. GAPDH used as reference gene for normalization of gene expression (n=3 donors).

**Supplemental Figure 5.**
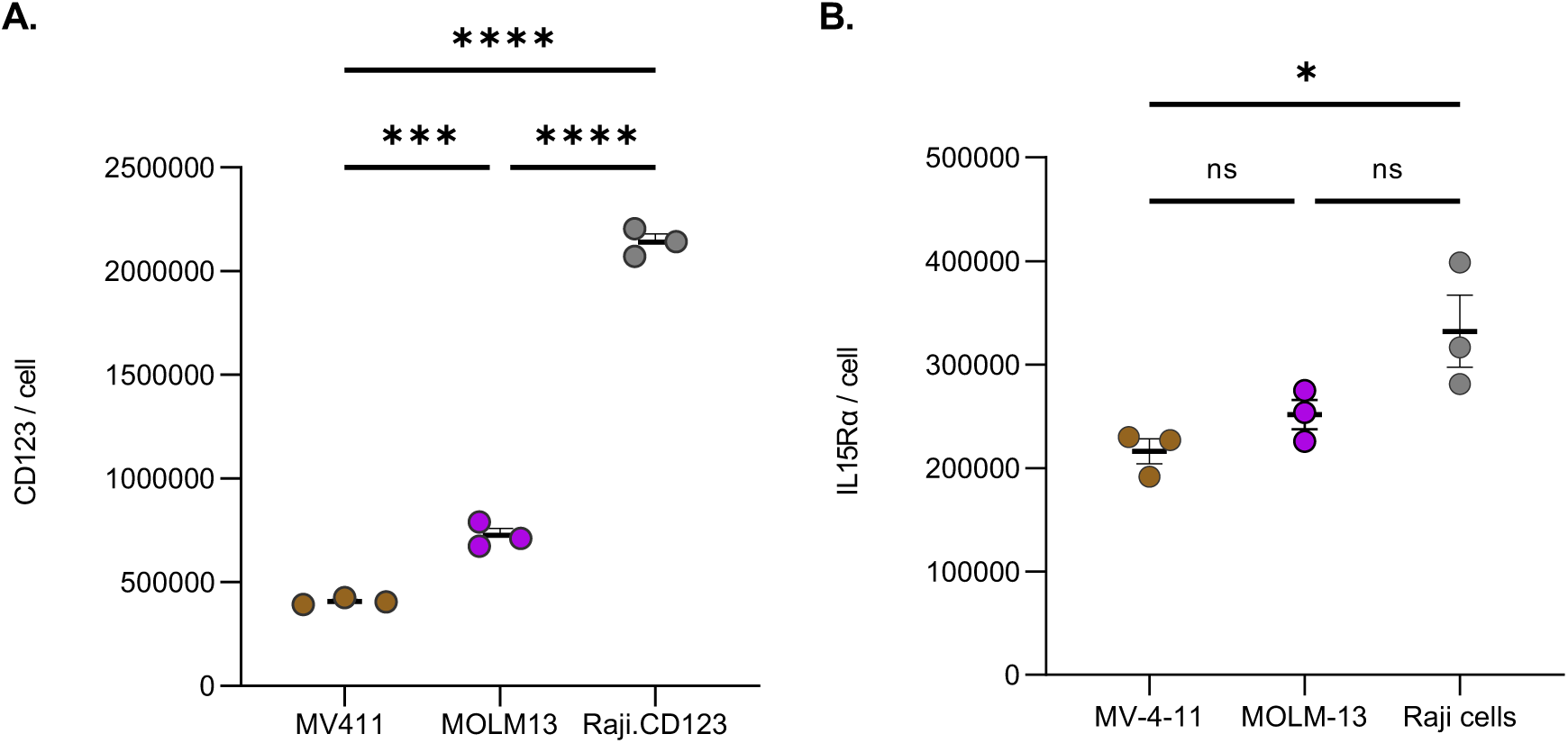
Surface antigen quantification on target cells. Quantification of (**A**) CD123 and (**B**) IL15Rα per cell (*p<0.05; ***p<0.001; ****p<0.0001).

**Supplemental Figure 6.**
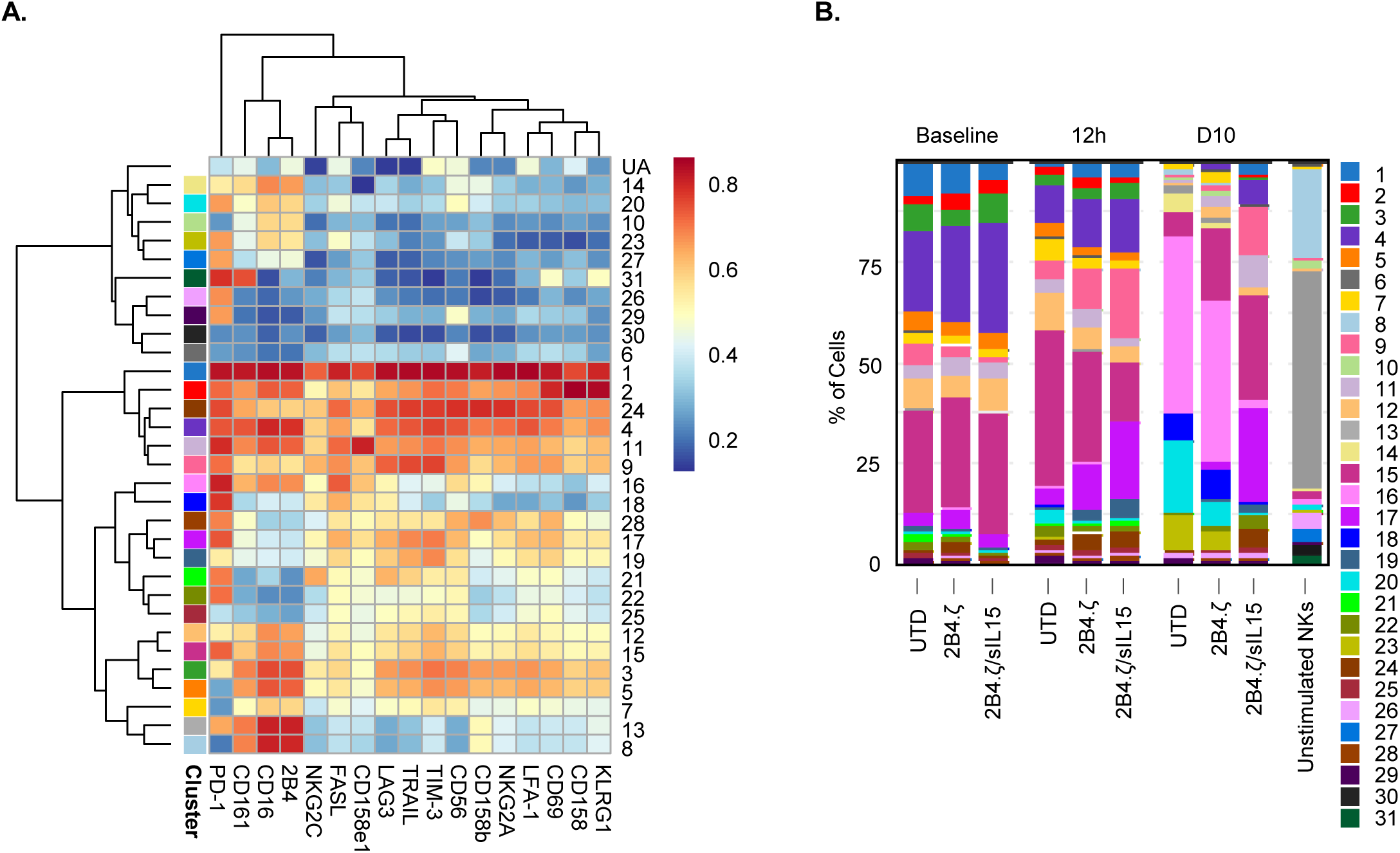
IL15-secreting CAR-NK cells demonstrate a distinct phenotype after chronic antigen stimulation. (**A**) Heatmap of flow cytometry data showing expression of 17 different NK cell surface markers. Heatmap coloring represents arcsinh transformed median marker intensities. (**B**) Bar plots of relative abundance of the 31 population subsets found in each sample. *The population clusters of the first panel (shown in Fig. 4) are not the same (numbers/colors) as the ones of the second panel, shown here.* Unstimulated, freshly isolated NK cells are used as controls.

**Supplemental Figure 7.**
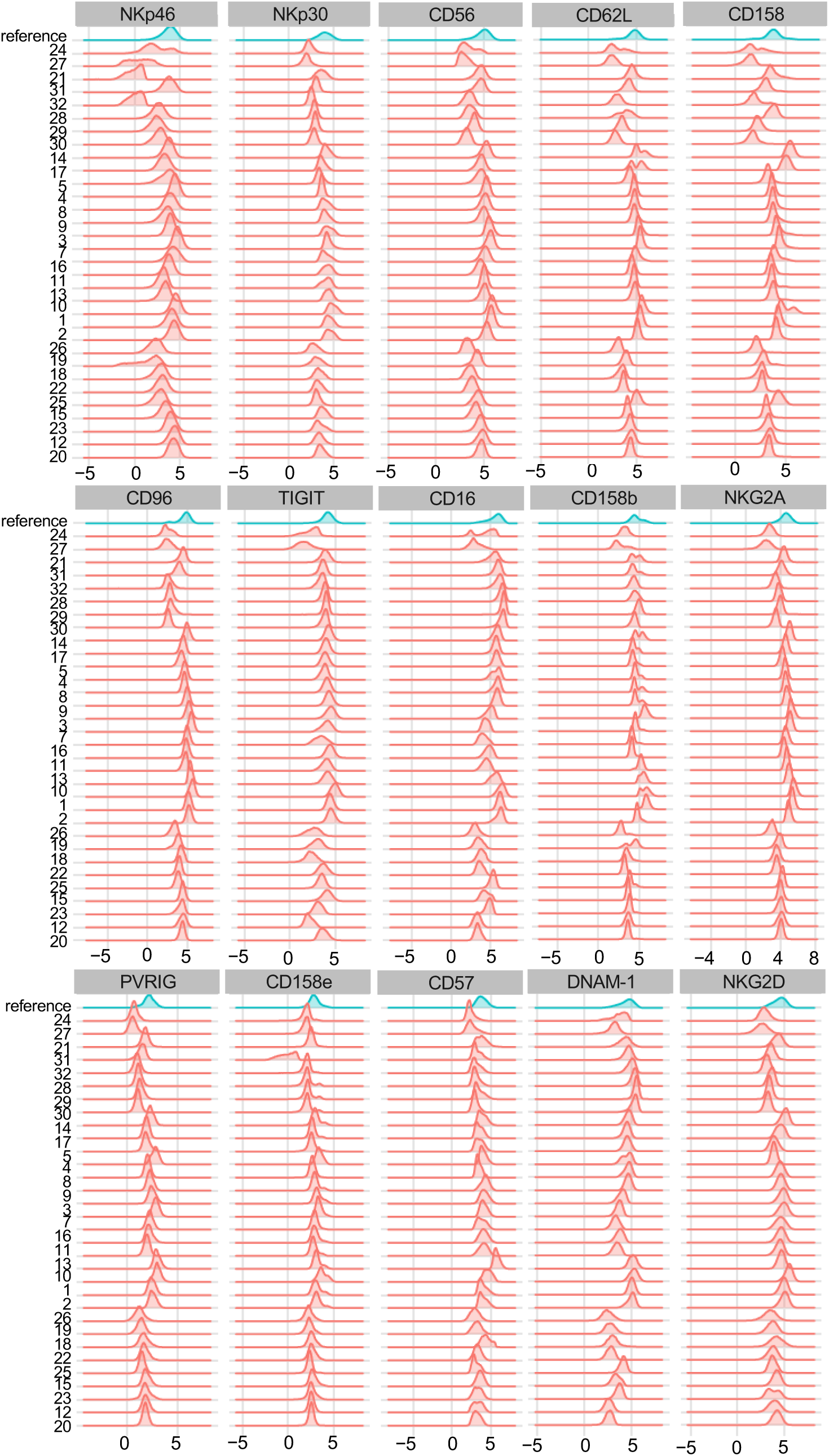
Distribution of marker intensities of indicated “Panel A” receptors in the identified 32 NK cell clusters. The NK cell clusters (on the left side) are named as in Figure 4. Red histograms represent the respective marker in each cluster. Blue histograms are a reference calculated from all cells.

**Supplemental Figure 8.**
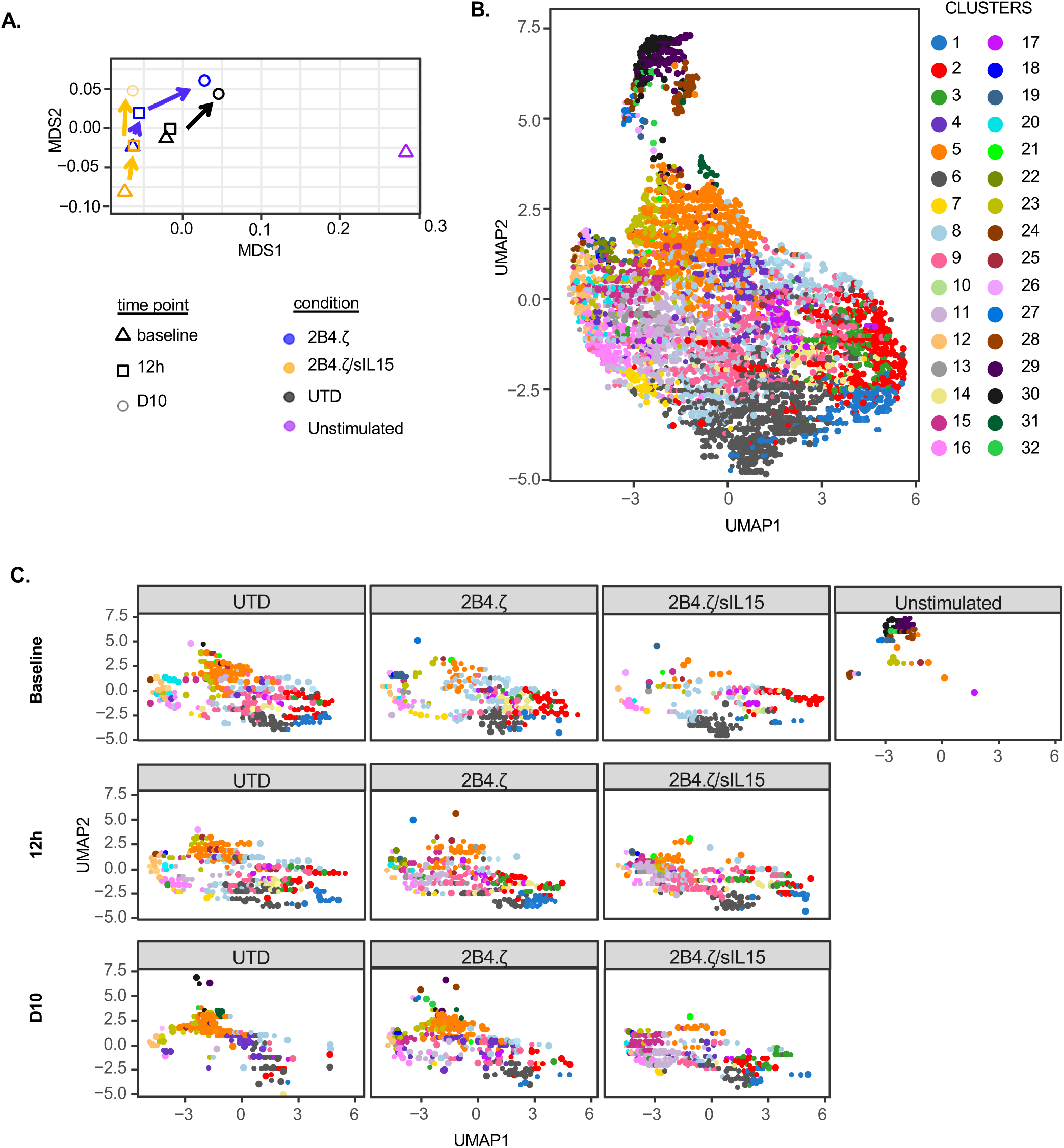
Visual representation of “Panel A” immunophenotype data. (**A**) Multidimensional scaling (MDS) plot. Unstimulated, freshly isolated NK cells are used as controls. Arrows indicate the transitions across timepoints of experiment. (**B**) Uniform Manifold Approximation and Projection (UMAP) plot was generated based on the arcsinh-transformed expression of the 15 expression markers in the NK cells from the whole dataset. Cells are colored according to the 32 clusters generated after manually merging the 40 meta-clusters obtained with FlowSOM. (**C**) Individual UMAP plots of different NK cell conditions. Different time points indicated on the left side (baseline, 12h, D10).

**Supplemental Figure 9.**
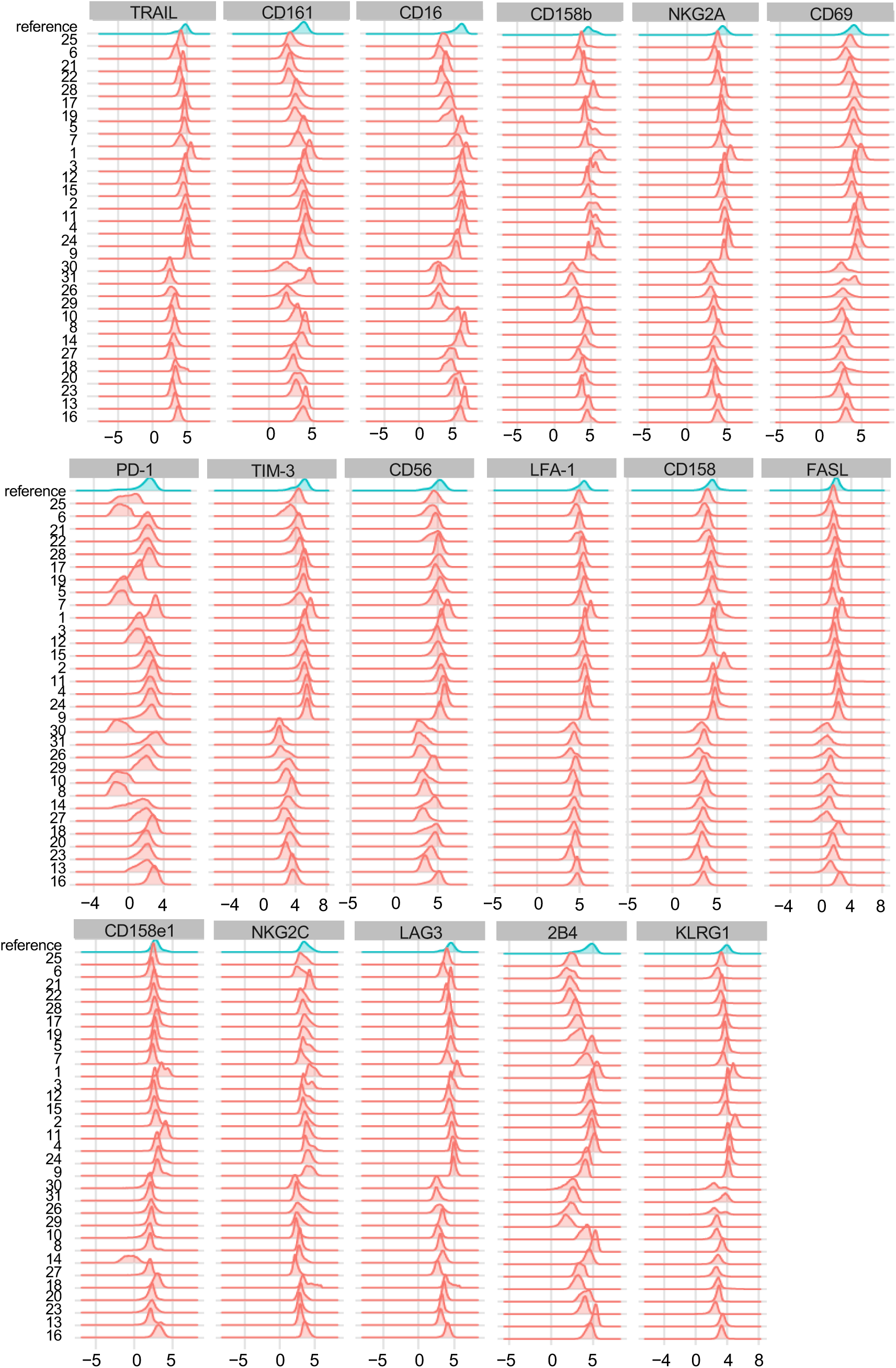
Distribution of marker intensities of the indicated “Panel B” receptors in the 31 NK cell clusters. The NK cell clusters (on the left side) are specific for Supplemental Fig. 6. Red histograms represent the respective marker in each cluster. Blue histograms are a reference calculated from all cells.

**Supplemental Figure 10.**
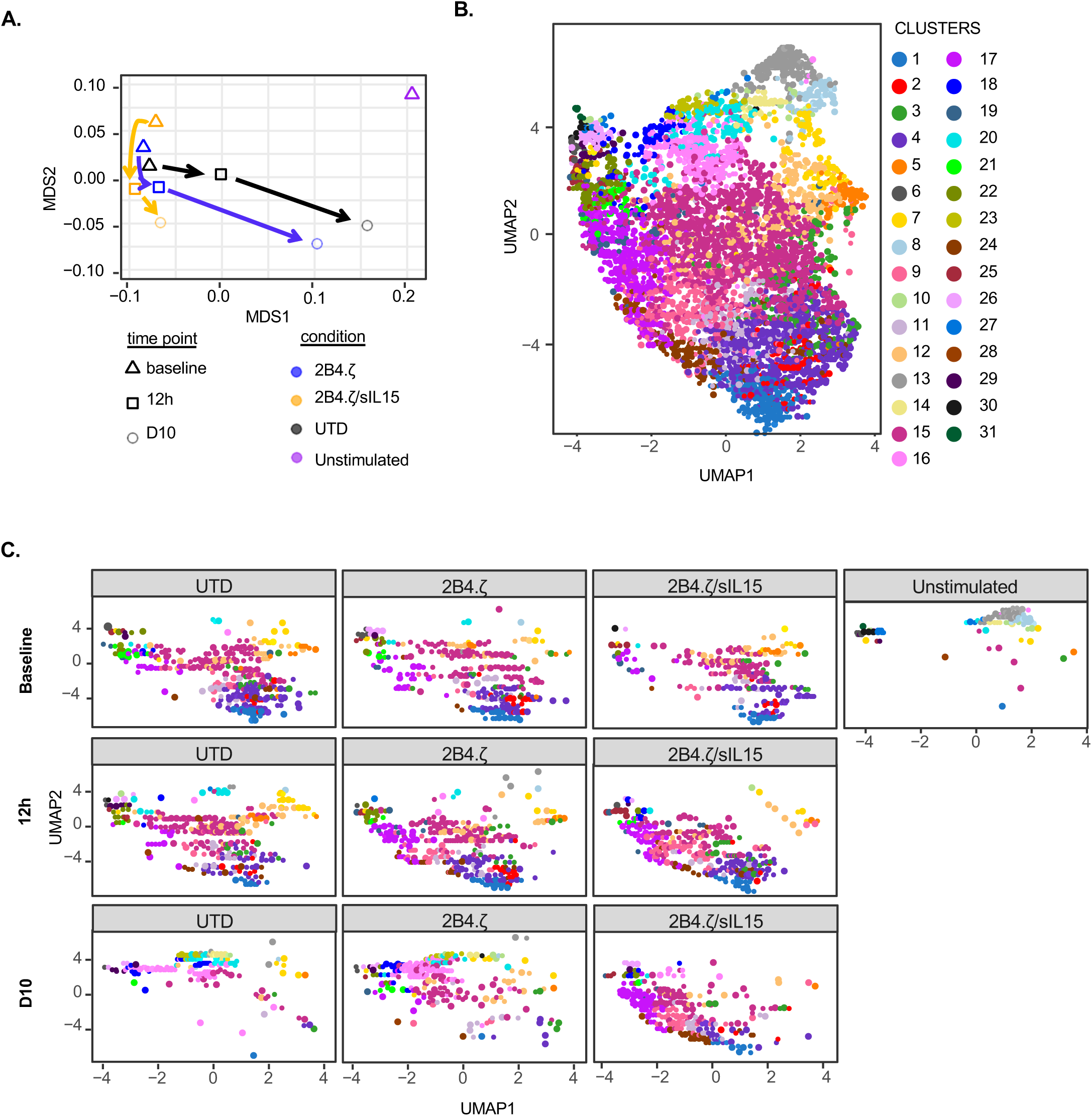
Visual representation of “Panel B” immunophenotype data. (**A**) Multidimensional scaling (MDS) plot. Unstimulated, freshly isolated NK cells are used as controls. Arrows indicate the transitions across timepoints of experiment. (**B**) Uniform Manifold Approximation and Projection (UMAP) plot was generated based on the arcsinh-transformed expression of the 17 expression markers in the NK cells from the whole dataset. Cells are colored according to the 31 clusters generated after manually merging the 40 meta-clusters obtained with FlowSOM. (**C**) Individual UMAP plots of different NK cell conditions. Different time points indicated on the left side (baseline, 12h, D10).

**Supplemental Figure 11.**
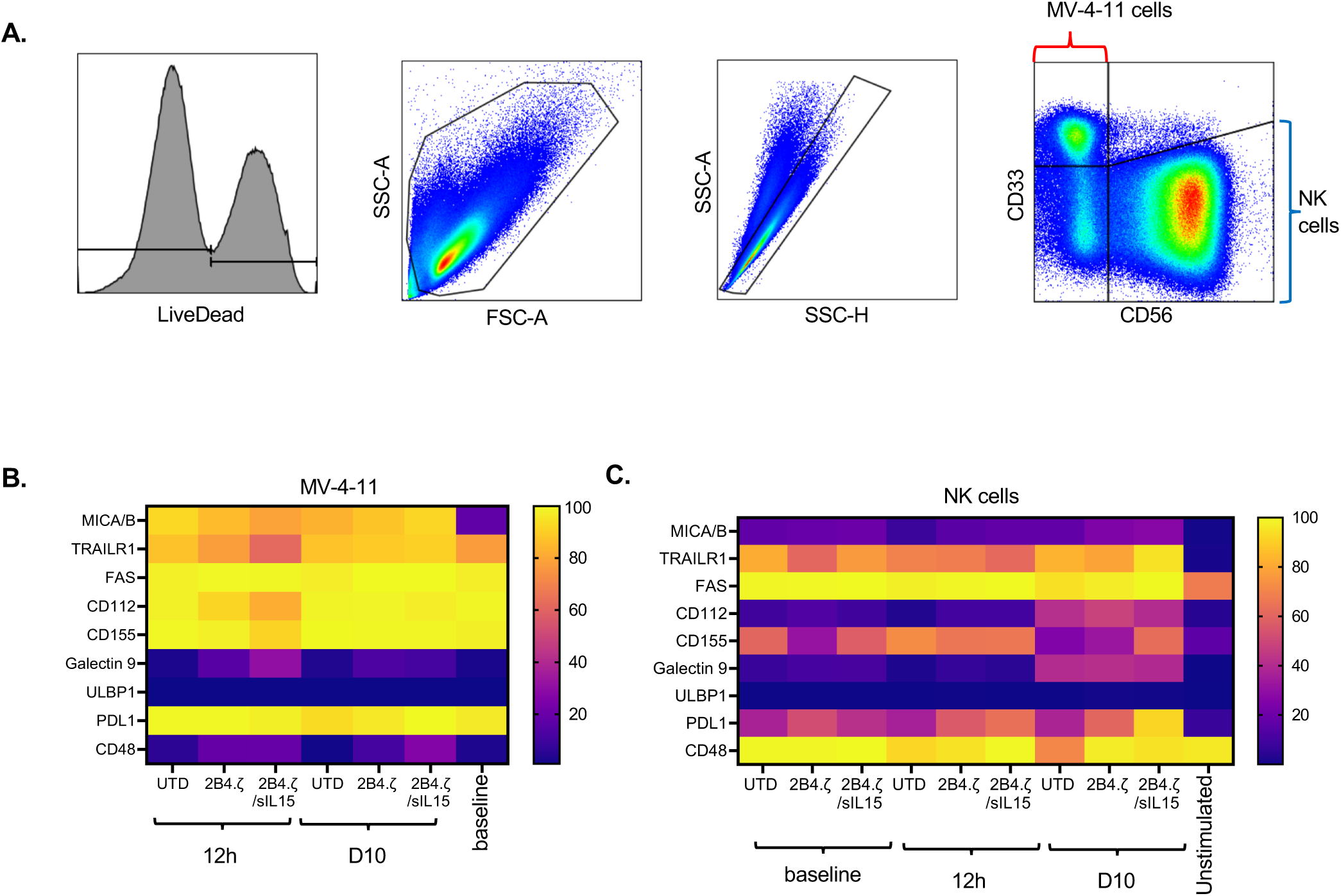
Expression of NK cell receptor ligands on target and NK cells. (**A**) Gating strategy. (**B**) Heatmap of the percent (%) expression of indicated ligands on MV-4-11 cells or (**C**) NK cells at each experimental time point.

**Supplemental Figure 12.**
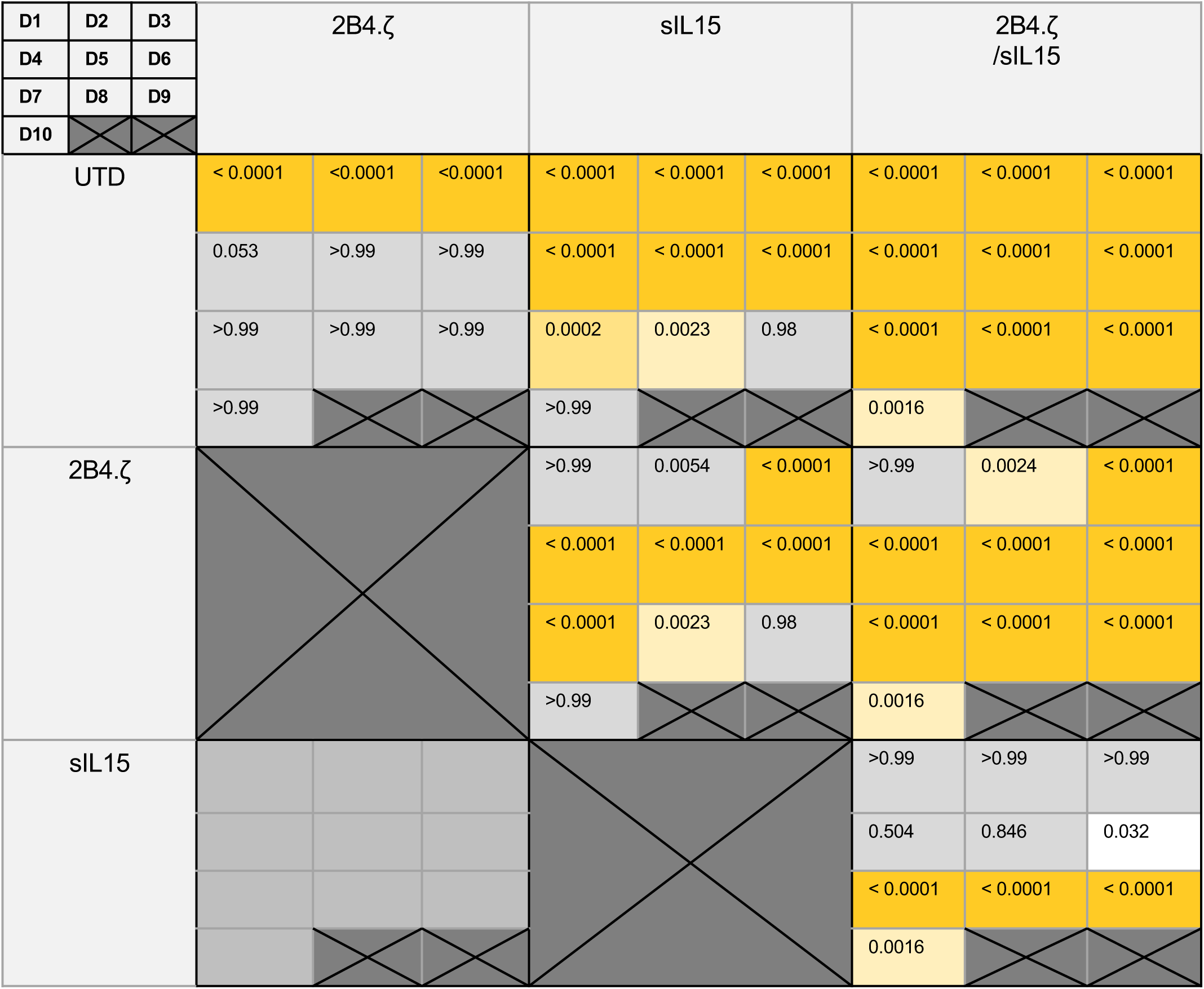
Comparisons of mean percent (%) cytotoxicity between different NK cell conditions in the serial stimulation assay shown in Fig. 4E. Each cell is subdivided to represent days 1-10 as indicated in upper left corner. Every cell’s value signifies the p-value of each comparison for every day. P values generated with ordinary 2-way ANOVA corrected for multiple comparisons using the method of Bonferroni.

**Supplemental Figure 13.**
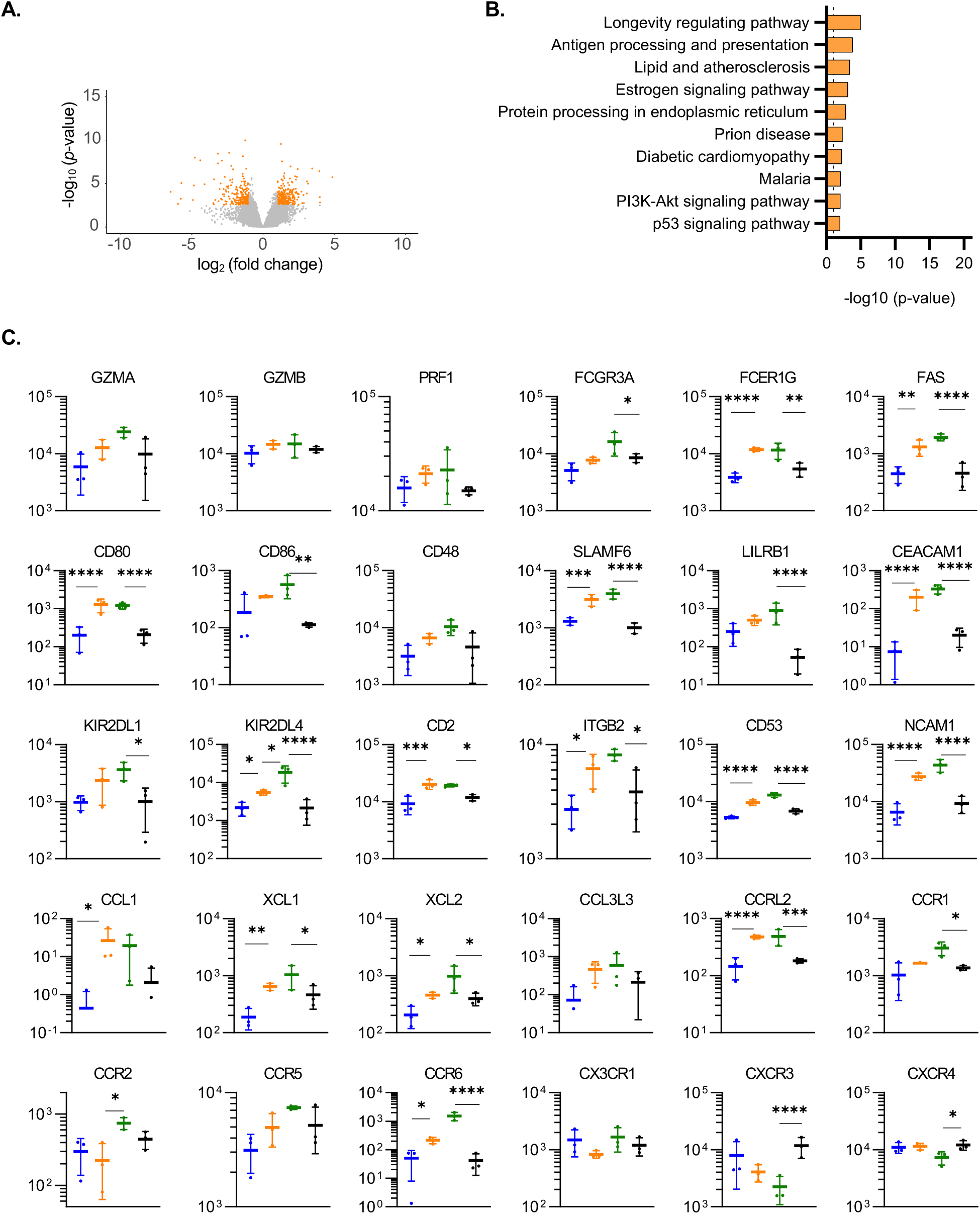
Transcriptomic evaluation of individual genes and differential expression analysis of IL15 secreting NKs. (**A**) Volcano plot representing upregulated and downregulated genes in 2B4.ζ/sIL15 compared to sIL15 NK cells. Orange dots on the left represent downregulated and dots on the right represent upregulated genes. Location of each data point is calculated as log2(FC) × −log10(p-value). Color cutoffs: p-value < 0.05; fold change cutoff >2 or <1/2. (**B**) Bar plot depicting the top 10 significant pathways enriched in the differentially expressed genes. KEGG 2021 Human gene Set Enrichment Analysis was used. Dotted line represents the cutoff p-value of 0.05.(**C**) Dot plots displaying the normalized expression of individual genes (n=3 independent donors). Statistical significance: *p<0.05; **p<0.01; ***p<0.001; ****p<0.0001.

**Supplemental figure 14.**
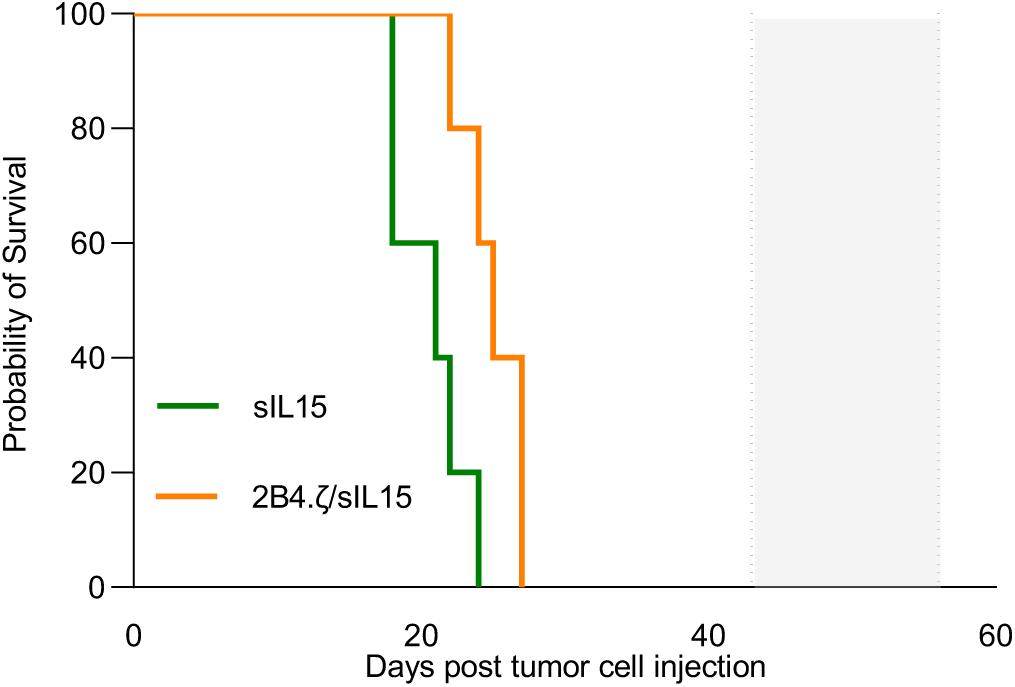
IL15 stimulated NK cells promote lethal toxicity of MV-4-11 engrafted mice. Kaplan–Meier survival analysis of MV-4-11 xenografts (n=5 mice each group). NSG mice received 1×10^6^ MV-4-11.ffLuc cells via tail vein injection on day 0. 10×10^6^ NK cells (Orange: 2B4.ζ/sIL15; Green: sIL15) were injected on D7. Dotted lines border grey shading representing usual survival window of MV-4-11 xenograft model.

**Supplemental Figure 15.**
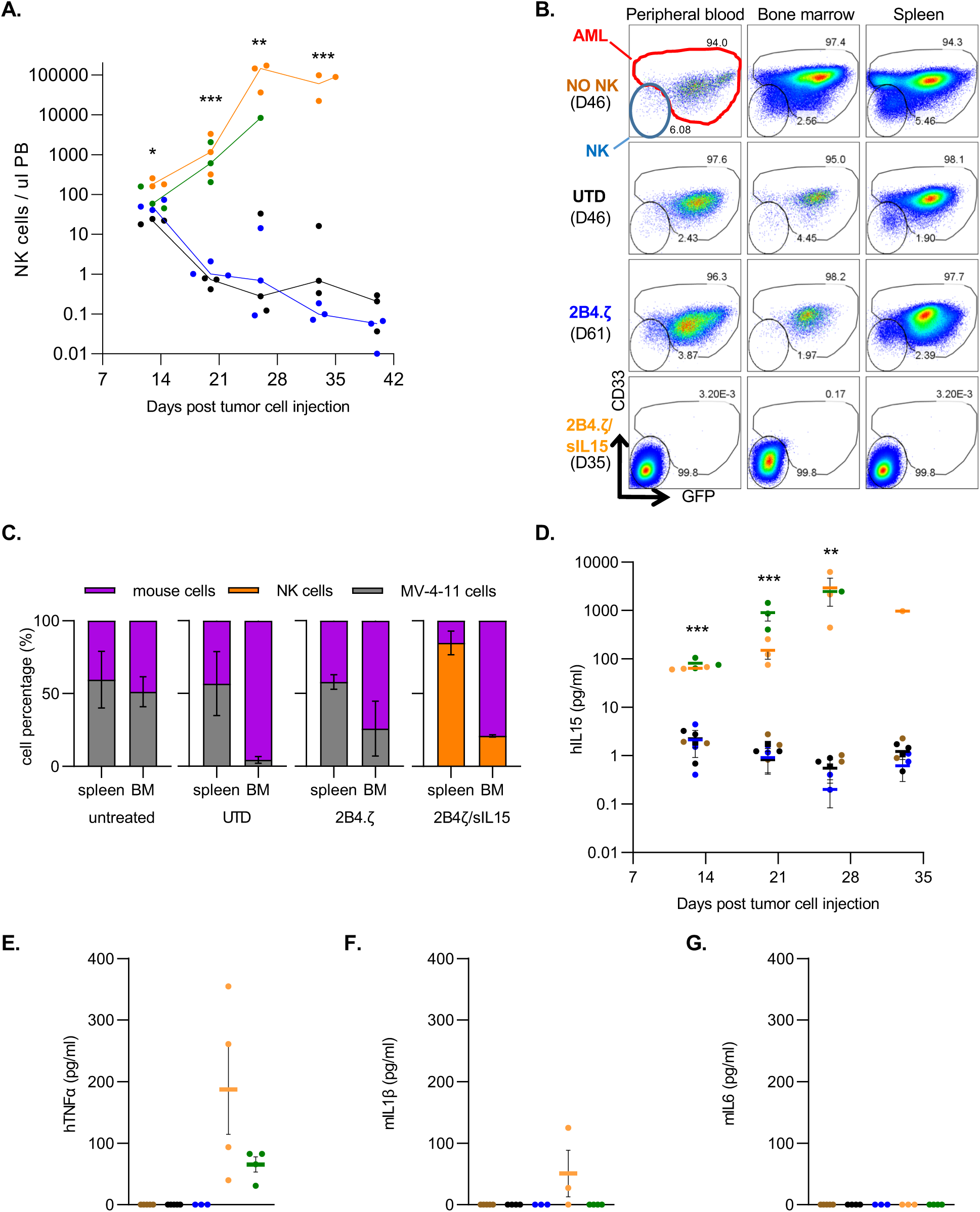
IL15 stimulation promotes NK cell expansion and inflammation *in vivo.* (**A**) Mouse peripheral blood (PB) was collected at indicated time points and analyzed via flow cytometry. NK cell numbers per microliter of mouse PB were tracked starting on day 13 of the experiment. Each dot represents cell numbers from a single mouse; line is at median. Asterisks indicate 2B4.ζ/sIL15 vs 2B4.ζ comparison. (**B**) FACS plots of hCD45(+) cells in the peripheral blood, bone marrow, and spleen of mice at necropsy. hCD45(+)CD33(-)GFP(-) cells: NK cells (blue). Percentage of cells populating NK or AML gates indicated. (**C**) Percentage (%) cell populations in spleen and bone marrow at necropsy (n=3 mice). Mean +/- SEM. (**D**) Human IL15 from peripheral blood of MV-4-11 engrafted mice drawn at the indicated time points was quantified (pg/mL) with ELISA. Asterisks indicate 2B4.ζ/sIL15 vs 2B4.ζ comparison. (**E**) human TNFα, (**F**) mouse IL1β and (**G**) mouse IL6 from peripheral blood of MV-4-11 engrafted mice drawn at necropsy and quantified with ELISA. Each dot derived from a single mouse. Mean +/- SEM. Statistical significance: *p<0.05; **p<0.01; ***p<0.001; ****p<0.0001.

**Supplemental Figure 16.**
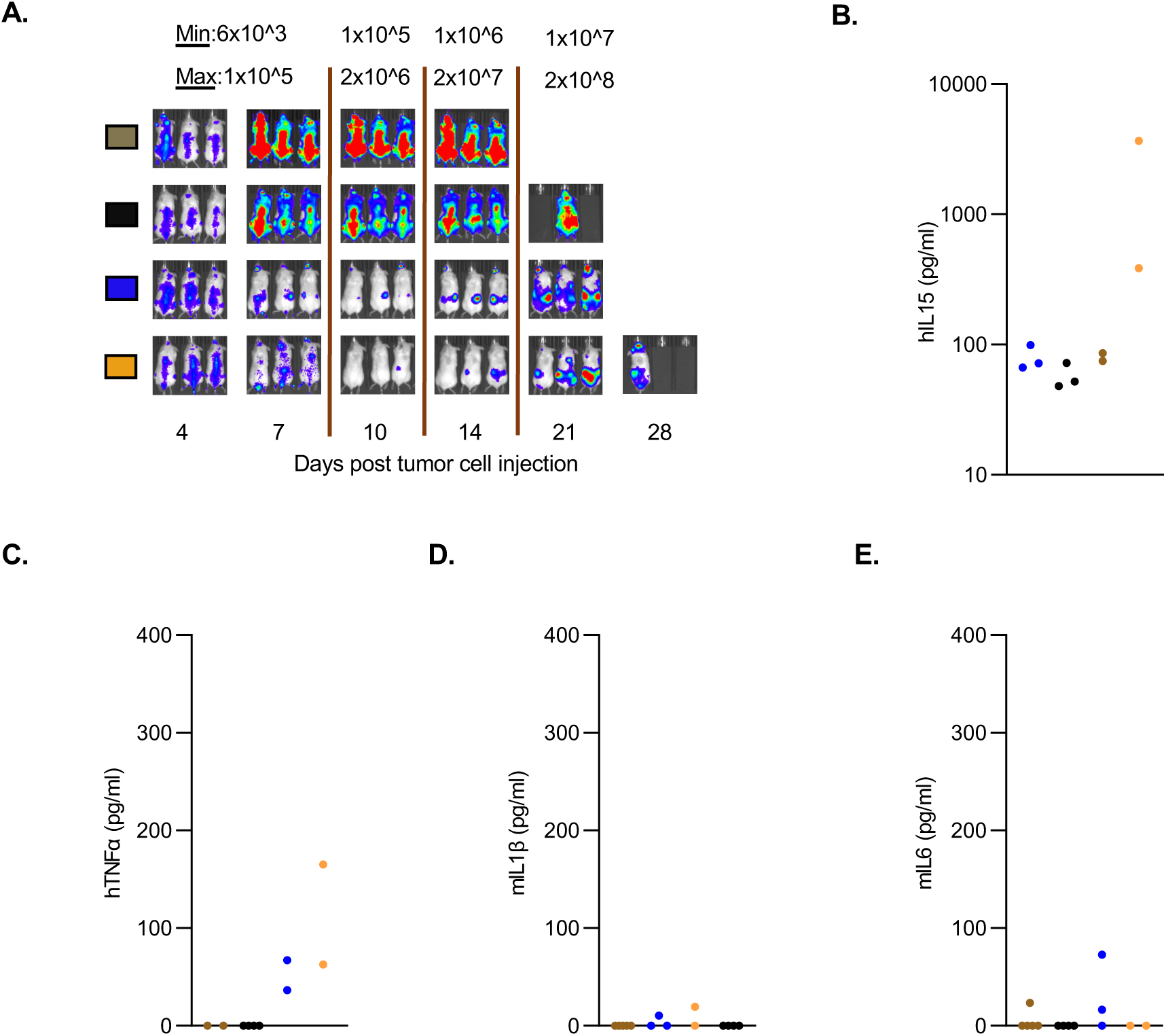
CAR-NK cell treatment has anti-tumor efficacy against MOLM-13 AML. (**A**) Mice were injected with 50,000 MOLM-13.ffLuc cells on day 0, followed by NK cell treatment in the following cohorts: No treatment (brown), 3 million unmodified (UTD) NK cells on days 4, 7, and 10 (black), 3 million 2B4.ζ NK cells on days 4, 7, and 10 (blue), 3 million 2B4.ζ*/*sIL15 NK cells on day 4 (orange). MOLM-13 proliferation was monitored with bioluminescence imaging (BLI) and was recorded as photons/sec/cm^2^/sr. Representative images of 3 mice per condition. The minimum and maximum values of the color scale used are defined.[min-max] (**B**) Human IL15, (**C**) human TNFα, (**D**) mouse IL1β and (**E**) mouse IL6 analysis from the peripheral blood drawn at necropsy from MOLM-13 engrafted mice. Cytokine measurement (pg/mL) with ELISA. Each dot represents data from a single mouse. Mean +/- SEM. n=2-5 mice. Statistical significance: *p<0.05; **p<0.01; ***p<0.001; ****p<0.0001.

## Supplemental Methods

### Cell lines

HEK293T (human embryonic kidney) , Raji (Burkitt’s lymphoma) and MV-4-11 (myelomonocytic leukemia) cell lines were purchased from the American Type Culture Collection (ATCC, Manassas, VA) and cultured in Dulbecco’s Modified Eagle’s Medium (DMEM; ThermoFisher Scientific Waltham, MA; 293T), Roswell Park Memorial Institute (RPMI; ThermoFisher Scientific; Raji) or Iscove’s Modified Dulbecco’s Medium (IMDM; ThermoFisher Scientific; MV-4-11) supplemented with 10% fetal bovine serum (FBS; HyClone, Logan, UT). MOLM-13 cell line was purchased from the Leibniz Institute (DSMZ, German Collection of Microoganisms and Cell Cultures) and cultured in RPMI supplemented with 10% FBS. CD123 expressing Raji cells (Raji.CD123) were created by first subcloning the full length human CD123 coding sequence into a pCDH lentiviral backbone. Vesicular stomatitis virus G glycoprotein (VSV-G) pseudotyped lentiviral particles were produced using the pPACKH1 HIV Lentivector Packaging Kit (System Biosciences, Palo Alto, CA) according to the manufacturer’s instructions and used for Raji cell modification. CD123-positive cells were isolated using fluorescence-activated cell sorting (FACS) and antigen surface expression verified prior to use. All cells used for BLI-based cytotoxicity assays and/or our xenograft models were transduced with a retroviral vector carrying an enhanced green fluorescent protein (GFP) firefly luciferase fusion gene (GFP.ffLuc).^1^ GFP-positive cells were sorted and maintained in the appropriate culture medium. Luciferase expression was confirmed using D-luciferin and quantification of bioluminescence. All cells were cultured in a humidified atmosphere containing 5% CO2 at 37°C.

### Determination of Vector Copy Number (VCN)

Primer/probe-FAM was designed to the MMLV-derived psi present in pSFG and purchased from ThermoFisher Scientific. RNAseP primer/probe-VIC/TAMRA mix (Applied Biosystems #4403326) was used as comparison. Genomic DNA was isolated from CAR-NK cells and 25 ng used for amplification with TaqMan Universal PCR Mastermix (ThermoFisher) and the above primer/probe mixes on a C1000 Touch Thermal Cycler (Bio-Rad, Hercules, CA). The following amplification conditions were used: 50°C for 2 minutes, 95°C for 10 minutes, 40 cycles of 95°C for 15 seconds, 60°C for 1 minute. No-template, unmodified NK cells and a condition containing only plasmid were used as controls. Vector copy number calculation was performed using the 2^-ΔCt^ method.^2^

### Cytotoxicity assay

Bioluminescence (BL) based: NK cells were co-cultured with target cells expressing ffLuc at the indicated E:T ratios. D-luciferin was added to plate and BL measured per well. Mean percentage of specific lysis of triplicate samples was calculated as 100*(spontaneous death–experimental death)/(spontaneous death-background). Spontaneous death was measured with control wells containing only target cells. Flow cytometric: NK cells were cultured with target cells. NK and target cell numbers were measured using flow cytometric analysis and NK or target-cell specific markers as above, with dead cell exclusion.

### Cytokine secretion assay

100,000 NK cells were plated with an equivalent number of target cells in 0.2 mL media and cultured for 24 hours. Supernatant was collected and IFNγ quantification performed via ELISA (R&D Systems, Minneapolis, MT). For measurement of IL15 secretion, 1 million NK cells were plated in 2 mL media. After 24 hours, supernatant was collected and cytokine quantification was performed with ELISA (R&D Systems) according to the manufacturer’s instruction.

### Library preparation and RNA sequencing

For RNA sequencing experiments, RNA samples were converted to double stranded cDNA using the Ovation RNA-Seq System v2.0 kit (Tecan, Männedorf, Switzerland), which utilizes a proprietary strand displacement technology for linear amplification of mRNA without rRNA/tRNA depletion as per the manufacturer’s recommendations. This approach does not retain strand specific information. Quality and quantity of the resulting cDNA was monitored using the Bioanalyzer High Sensitivity kit (Agilent) which yielded a characteristic smear of cDNA molecules ranging in size from 500 to 2000 nucleotides in length. After shearing 500 nanograms of cDNA to an average size of 250 nucleotides with the Covaris S4 (Covaris Inc., Woburn, MA) library construction was completed with the Truseq Nano kit (Illumina; San Diego, CA) according to the manufacturer’s instructions. mRNA libraries were sequenced on an Illumina Novaseq 6000 instrument using 150bp paired-end dual indexed reads and 1% of PhiX control. Reads were aligned to GRCh38 using rsem version 1.3.0 with the following options – star-calc-ci-star-output-genome-bam-forward-prob 0.5.

---

**Supplemental Table 1.** Detailed list of the fluorophore-conjugated antibodies used in flow cytometry experiments.

| Marker | Fluorophore | Clone | Company |
| --- | --- | --- | --- |
| CD56 | BV421 | HCD56 | BioLegend |
| CD56 | APC | HCD56 | BioLegend |
| CD56 | BUV805 | NCAM16.2 | BD |
| CD16 | BUV563 | 3G8 | BD |
| CD33 | PerCP/Cyanine5.5 | P67.6 | BioLegend |
| CD33 | BV785 | WM53 | BioLegend |
| His-tag | APC | J095G46 | BioLegend |
| His-tag | PE | J095G46 | BioLegend |
| CD158b (KIR2DL2/2DL3/2DS2) | BUV661 | CH-L | BD |
| CD158e1 (KIR3DL1) | BV480 | DX9 | BD |
| CD158 (KIR2DL1/S1/S3/S5) | APC/Fire™ 750 | HP-MA4 | BioLegend |
| NKG2A (CD159a) | UV737 | 131411 | BD |
| NKp30 (CD337, NCR3) | BB700 | p30-15 | BD |
| NKp46 (CD335 , NCR1) | BB515 | 9E2/NKp46 | BD |
| CD57 | BV605 | QA17A04 | BioLegend |
| CD62L | Red 718 | SK11 | BD |
| PVRIG | Alexa Fluor 405 | 2334A | R&D Systems |
| TIGIT | BUV395 | 741182 | BD |
| CD96 | BV711 | 6F9 | BD |
| NKG2D (CD314) | PE-Cy7 | 1D11 | BioLegend |
| DNAM-1 (CD226) | PE/Dazzle™ 594 | 11A8 | BioLegend |
| TRAIL (CD253) | BV711 | RIK-2 | BD |
| FAS-L ( CD178) | BV421 | NOK-1 | BioLegend |
| TIM-3 (HAVCR2) | BB700 | 7D3 | BD |
| LAG3 (CD223 ) | BV650 | 11C3C65 | BioLegend |
| PD-1 (CD279) | BB515 | EH12.1 | BD |
| NKG2C (CD159c) | BV605 | 134591 | BD |
| 2B4 (CD244) | PE-CF594 | 2-69 | BD |
| KLRG1 (MAFA) | PE-Cy7 | 2F1/KLRG1 | BioLegend |
| CD69 | BUV805 | FN50 | BD |
| LFA-1 (CD11a) | Red 718 | G-25.2 | BD |
| CD161 | BUV395 | HP-3G10 | BD |
| TRAIL-R1 (CD261) | BV421 | S35-934 | BD |
| FAS (CD95) | BB515 | DX2 | BD |
| MICA/MICB (PERB11) | PE-Cy7 | 6D4 | BioLegend |
| ULBP-1 | APC | 170818 | R&D Systems |
| CD48 | PE-CF594 | TÜ145 | BD |
| CD112 (Nectin-2) | PerCP/Cyanine5.5 | TX31 | BioLegend |
| CD155 (PVR) | V711 | TX24 | BD |
| galectin-9 | PE | 9M1-3 | BD |
| PD-L1 (CD274) | UV395 | MIH1 | BD |
| CD19 | PerCP/Cyanine5.5 | SJ25C1 | BioLegend |

**Supplemental Table 2.**
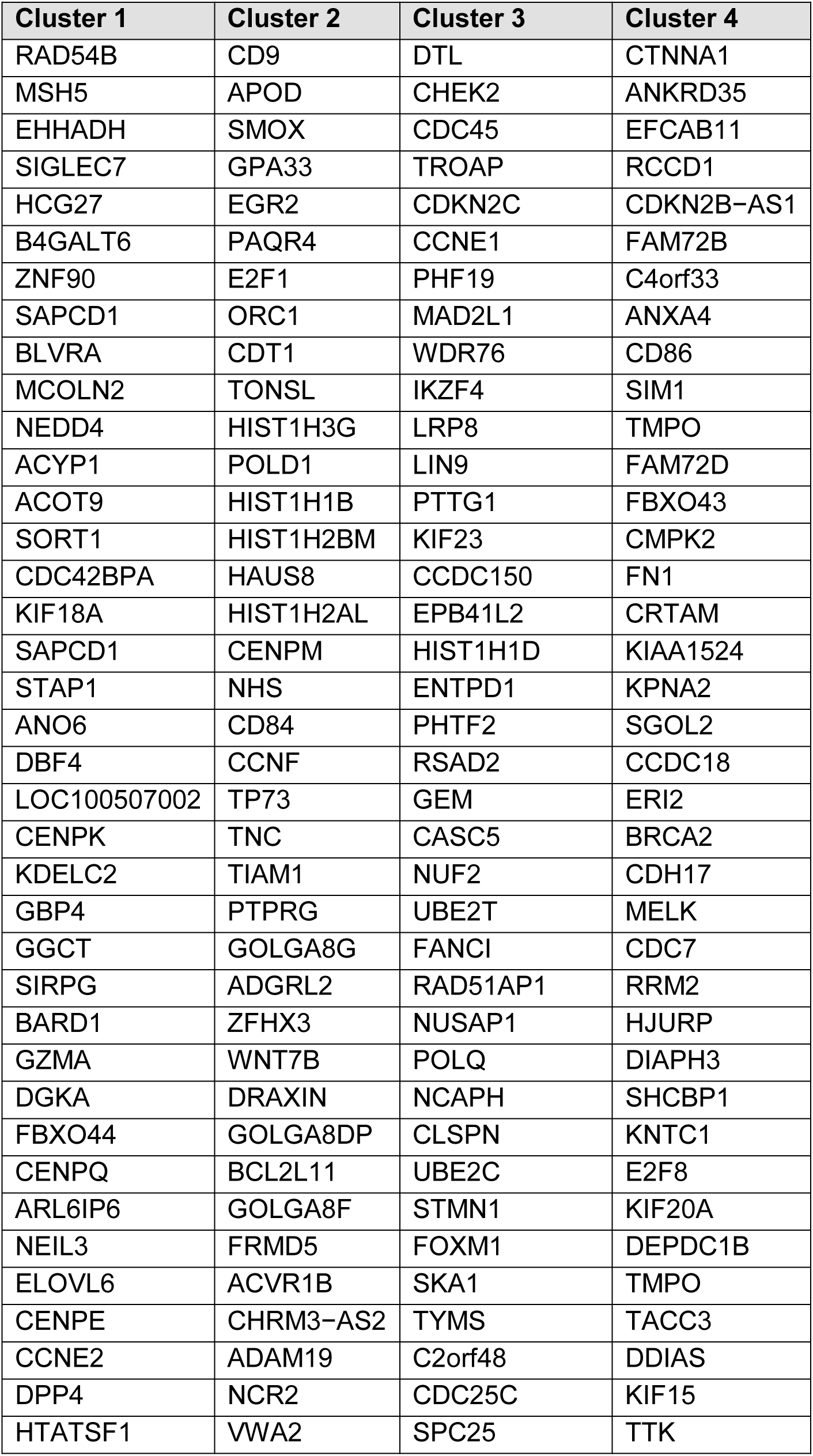
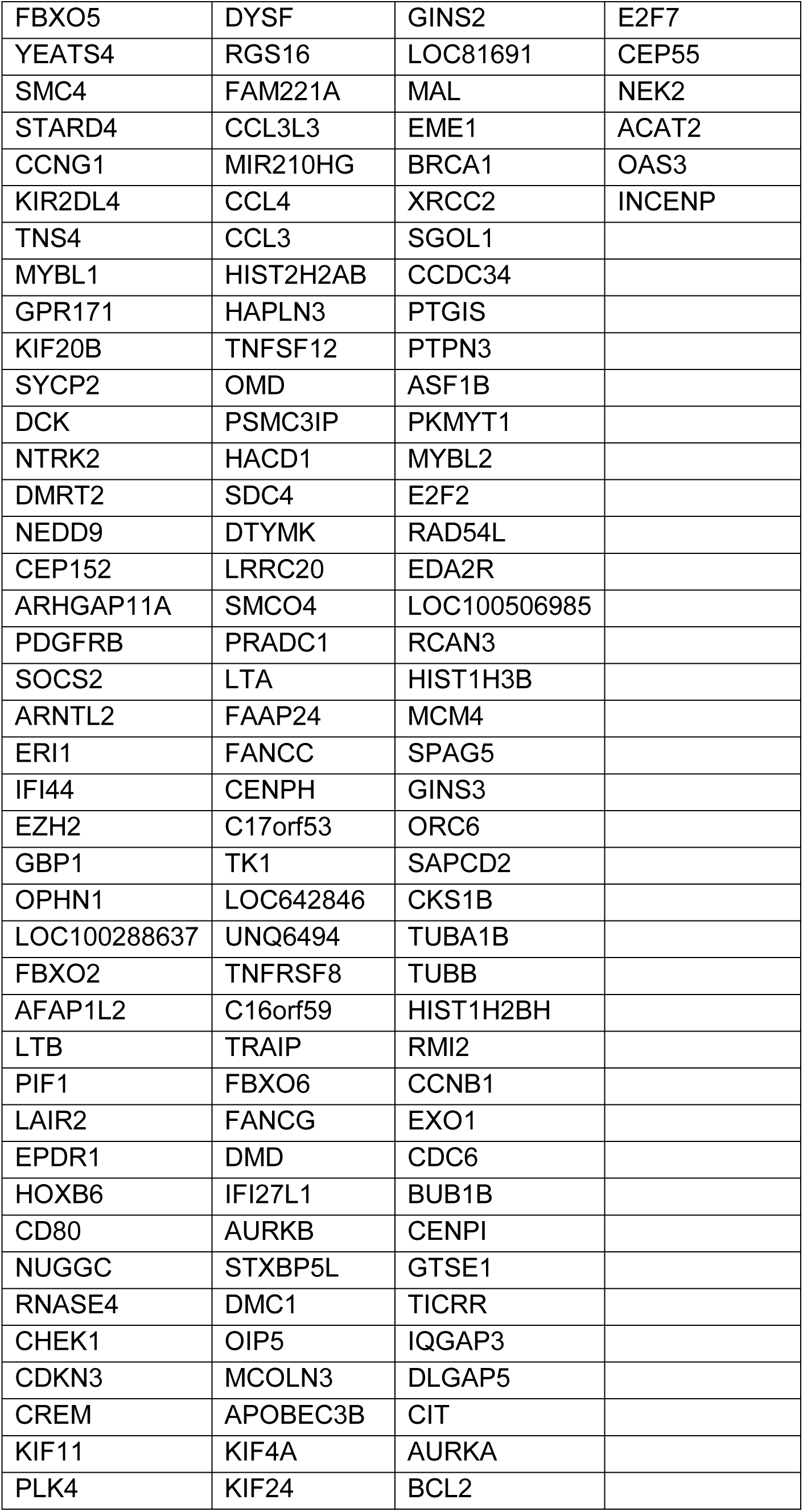
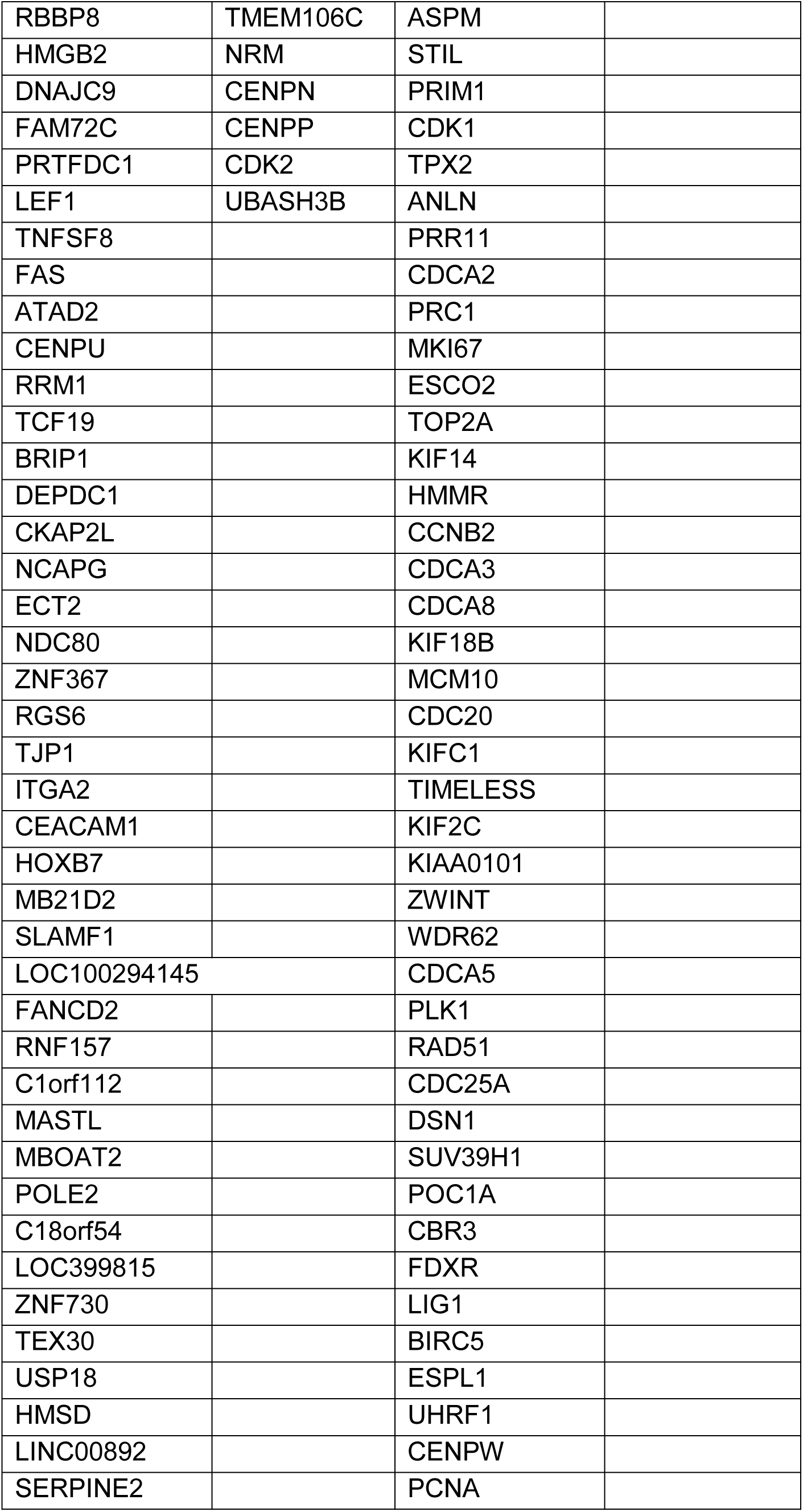
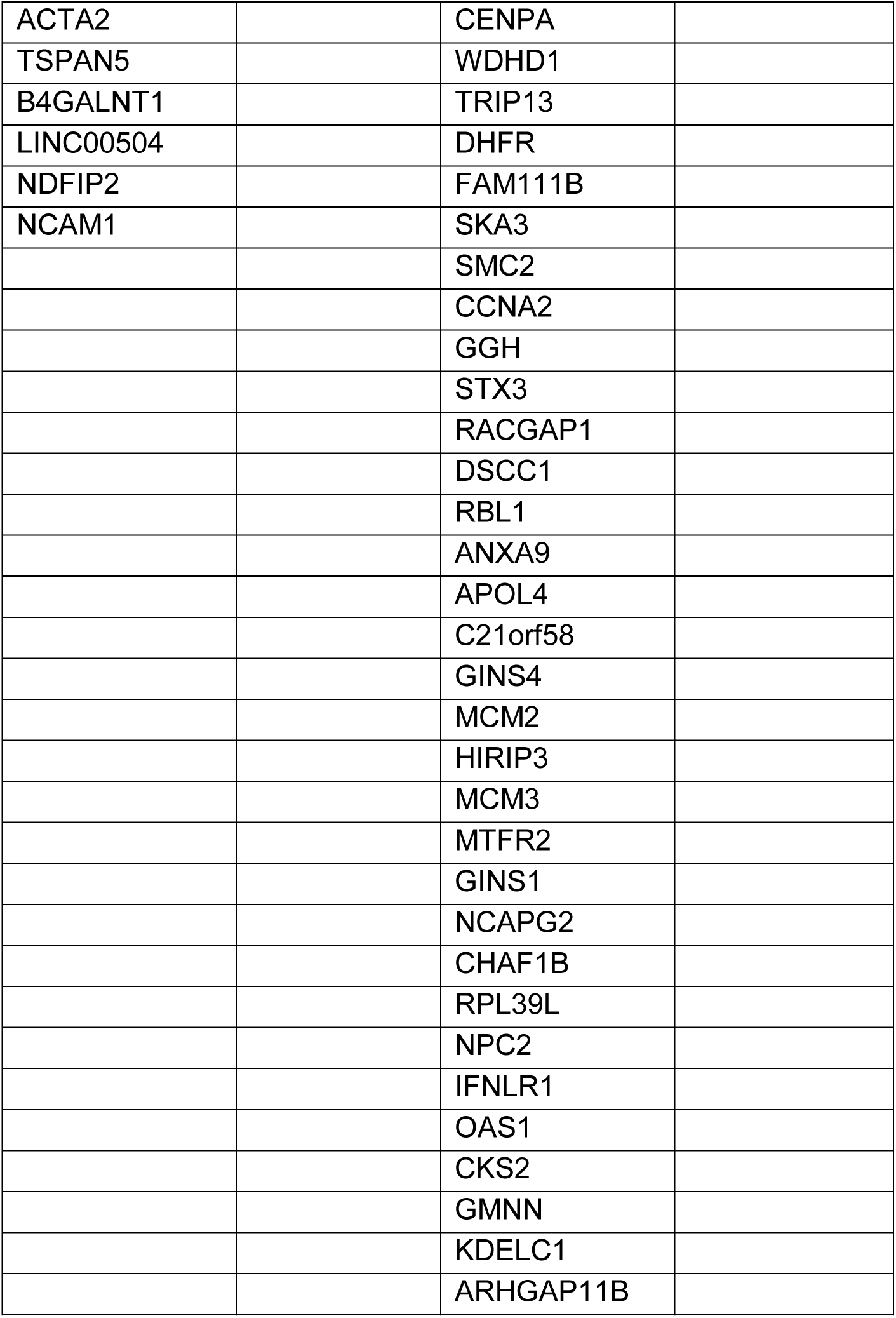
Genes included in the clusters 1-4 shown in Fig. 5D (listed from top to bottom of the cluster).

